# Conformational plasticity of human acid-sensing ion channel 1a

**DOI:** 10.1101/2024.12.11.628012

**Authors:** James Cahill, Kimberly A. Hartfield, Stephanie Andrea Heusser, Mette Homann Poulsen, Craig Yoshioka, Stephan Alexander Pless, Isabelle Baconguis

**Affiliations:** Vollum Institute, Oregon Health & Science University, Portland, OR 97239; University of Copenhagen, Department of Drug Design and Pharmacology, Jagtvej 160, 2100 Copenhagen, Denmark; Department of Biomedical Engineering, Oregon Health & Science University, Portland, OR 97239

**Author notes:** These authors contributed equally (James Cahill, Kimberly A. Hartfield). Correspondence: Isabelle Baconguis.

## Abstract

Acid-sensing ion channels (ASICs) are typically activated by acidic environments and contribute to nociception and synaptic plasticity. ASIC1a is the most abundant subunit in the central nervous system and forms homomeric channels permeable to Na^+^ and Ca^2+^, making it a compelling therapeutic target for acidotic pathologies including stroke and traumatic brain injury. However, a complete conformational library of human ASIC1a in its various functional states has yet to be described. Using cryo-EM, we obtained hASIC1a structures across a pH range between 8.5 and 5.7, as well as in the presence of a toxin agonist and a gating-modulating mutation. We identify six major conformations that establish linear transmembrane helices to be associated with an open state, delineate mechanistic differences between proton and toxin activation, and demonstrate that desensitization leads to unexpected conformational diversity in the transmembrane domain. Together, they provide a three-dimensional rationalization of decades of structure-function studies on ASIC.

## Introduction

Acid-sensing ion channels (ASICs) are trimeric proton-gated sodium channels. ASICs are members of the epithelial sodium channel (ENaC)/degenerin family of ion channels, and are widely expressed, including in the central and peripheral nervous system. They are involved in important physiological processes such as synaptic transmission^1^, memory^2^ and fear response^3^. ASIC activation is also implicated in neuronal injury during ischemic stroke^4,5^ and myocardial ischemia-reperfusion injury due to cardiac infarction^6^, and plays a role in pain^7^ and other pathologies, such as epilepsy, anxiety and addiction^8^.

Four genes (*ASIC1-4*) express six ASIC isoforms (ASIC1a, 1b, 2a, 2b, 3, and 4), forming functional homo- and heterotrimers^9^. The identified isoforms can all transition between resting closed, activated open, and desensitized closed states, with pH sensitivity, desensitization kinetics, and ion selectivity differentially affected by subunit composition. ASIC1a activation typically has a pH_50_ of 6.5-6.7 and desensitizes within tens to hundreds of milliseconds.

Structurally, each protomer in a trimer has a short intracellular N-terminus that directly precedes the first transmembrane helix (TM1), part of which can form a membrane-embedded re-entrant loop^10–12^. This pre-TM1 region also contains a conserved HG (His-Gly) motif. TM1 is followed by a large extracellular domain (ECD) that is responsible for proton sensing, and contains a set of β-strands that form a so-called “molecular clutch” controlling channel desensitization^13^. The ion-conducting channel pore is lined by the second transmembrane helix (TM2). TM2 is situated downstream of the ECD and contains a GAS (Gly-Ala-Ser) motif that is highly conserved throughout the ENaC/degenerin family. The channel pore is primarily permeable to sodium and potassium, while calcium can both permeate and inhibit the channel. This permeability profile has possible clinical relevance due to the role of sodium and calcium influx in acidosis-associated brain injury during ischemic stroke^5^.

The potential for pharmacological modulation of ASIC function is apparent from the diversity of naturally occurring toxins targeting ASICs, primarily derived from the venoms of snake, tarantula, and sea anemone species with various activating, inhibiting, and modulating effects^14^. The binding modes of several such toxins are known from structural characterization using either chicken ASIC1^10,15,16^ or human ASIC1a^17^, but a full description of the ASIC-toxin interaction paradigm would benefit from improved understanding of the conformational dynamics across the relevant functional states^14^. For example, the toxin agonist MitTx from *Micrurus tener tener* has been proposed to interact primarily with the closed resting state to induce conformational rearrangements that lead to an open state^7,10^, and therefore interpretation of the mechanism of binding and channel activation would be limited without structures of both the closed resting state and MitTx-bound open state. Furthermore, it remains unclear whether the opening mechanism and resultant open state conformation are preserved between channels activated by protons and MitTx.

Understanding of both the physiological function of ASIC and the modulatory potential of pharmacological agents is benefitted by enhanced structural understanding of the conformational diversity of its resting, conducting, and desensitized states. However, past studies have presented more focused structural perspectives, thus falling short of providing a comprehensive compendium of ASIC1a structures that would allow unambiguous assignment of structural correlates to the different functional states. Using single particle cryo-electron microscopy and sampling a wide pH range within a single buffer chemistry, we report the structures of six distinct conformations of human ASIC1a. While we observe a three-step progression from closed to open to desensitized conformations in the extracellular domain (ECD), the transmembrane domain (TMD) presents a surprising conformational diversity, including three distinct TMD conformations associated with the desensitized ECD. This ECD- and TMD-specific structural diversity offers crucial insights into the diverse functional landscape of ASICs in the nervous system.

## Results

### hASIC1a structural characterization under multiple pH conditions

Structural characterization of ASIC1 at different pH conditions has previously relied on buffering systems that were individually selected for their buffering capacity at the pH of interest (e.g. Tris to observe a putative closed, resting state at higher pH^12,13,17,18^ or sodium acetate for probing structural changes at low pH^10,15,16^). In order to conduct structural studies across a broad pH range, while minimizing perturbation by buffer chemistries, we chose a sodium phosphate buffering system that has the ability to buffer solutions in a comparatively wide range. To validate the use of phosphate buffer for structural characterization of hASIC1, activation and steady-state desensitization (SSD) curves were determined in a simple buffer of phosphate and sodium chloride, and in a more physiological buffer (ND96; HEPES based) (Figures S1A-S1D). Both buffer systems resulted in pH-dependent responses in heterologously expressed hASIC1a in *Xenopus* oocytes and HEK cells, although the pH_50_ values in phosphate buffer were shifted in the alkaline direction for activation (ΔpH 0.26 and 0.53 pH units for *Xenopus* oocytes and HEK cells, respectively) and SSD (ΔpH 0.27 units for *Xenopus* oocytes) (Figure S1; Table S1). In HEK cells, we observed a small, sustained current induced by phosphate buffer at pH 7.5 (Figure S1E), providing functional evidence for the occurrence of open conformations at this pH.

The conformational landscape of hASIC1a was studied at four pH conditions using single particle cryo-EM: pH 8.5, 7.5, 6.5, and 5.7. Full-length hASIC1a was expressed in HEK293S cells, extracted, and purified with digitonin as the solubilizing and stabilizing detergent. By varying the pH, we solved eight structures of wild-type hASIC1a, giving rise to six different conformations (Figure 1; Figure S2; Tables S2-S4). The conformations were differentiated on the basis of several key structural elements summarized in Table 1. For clarity, we have labeled all our structures according to the functional state associated with their ECD conformation (**C**losed, **O**pen or **D**esensitized) and TM architecture (**D**omain **S**wapped or **L**inear) in subscript. To specify a particular structure observed at a pH condition, the corresponding pH value is added in the label.

**Figure 1:**
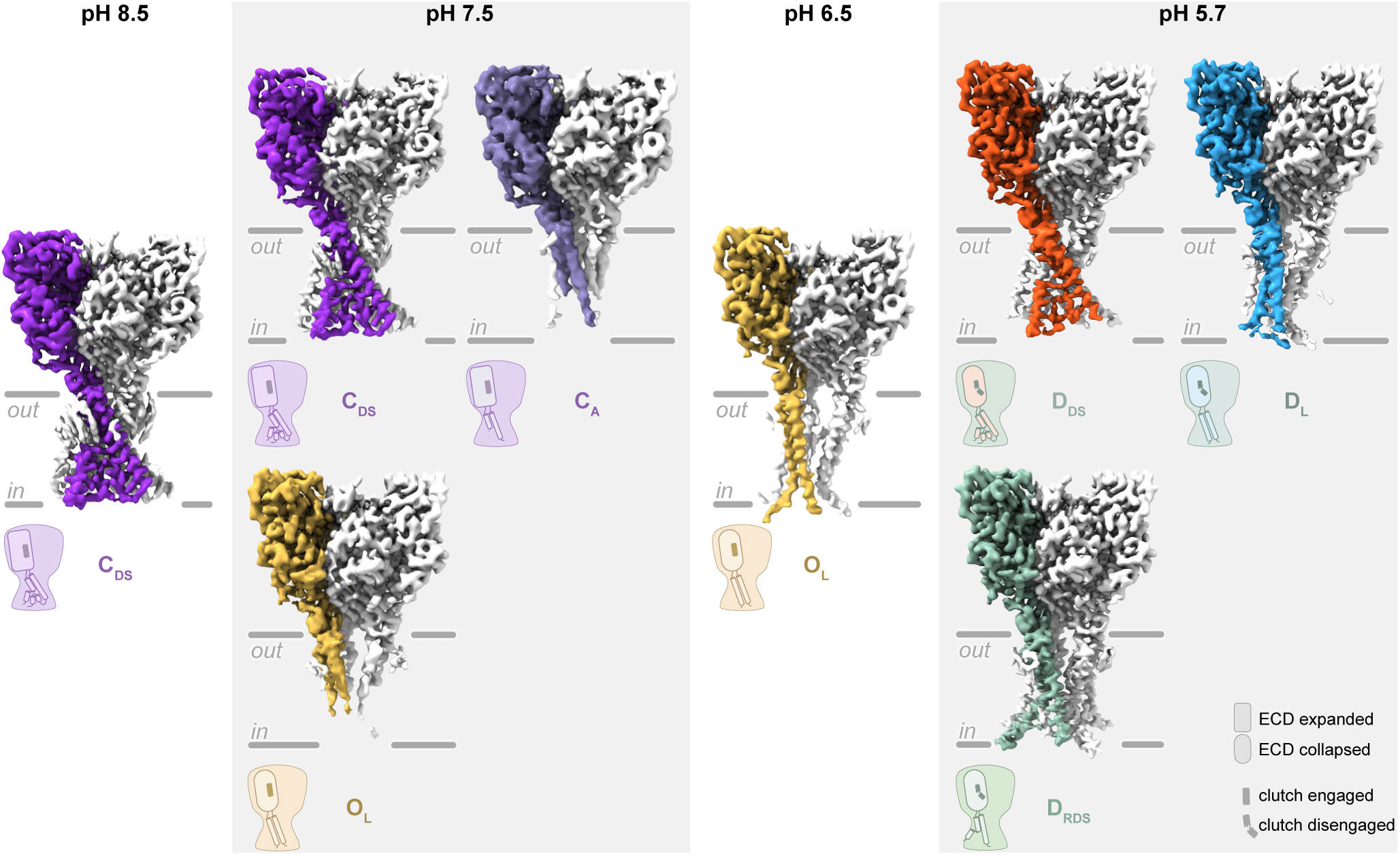
Conformational plasticity of hASIC1a by pH modulation. Maps of hASIC1a are arranged by the pH at which the samples were prepared. Map regions corresponding to an individual hASIC1a subunit are colored; colors are coded by conformation. The orientation and approximate position of the homotrimer within the plasma membrane is indicated with gray lines. Cartoon schematics are shown beneath the corresponding map and represent the major morphological features of TMD conformation, acidic pocket conformation, clutch orientation, and apparent role in the mechanistic cycle (color of background shape: purple for resting; gold for conducting; and green for desensitized). See also Figures S1-S3.

**Table 1.**
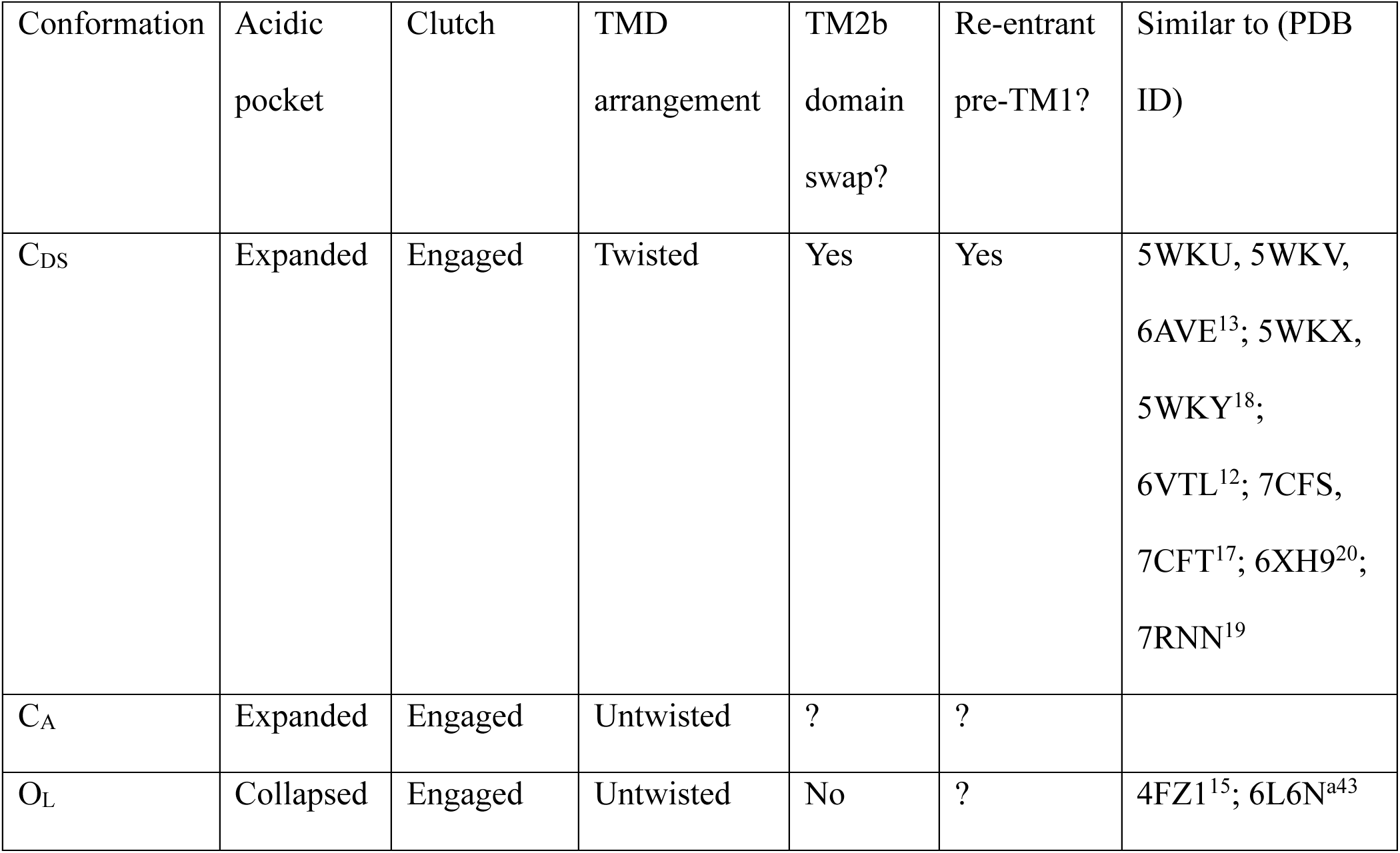
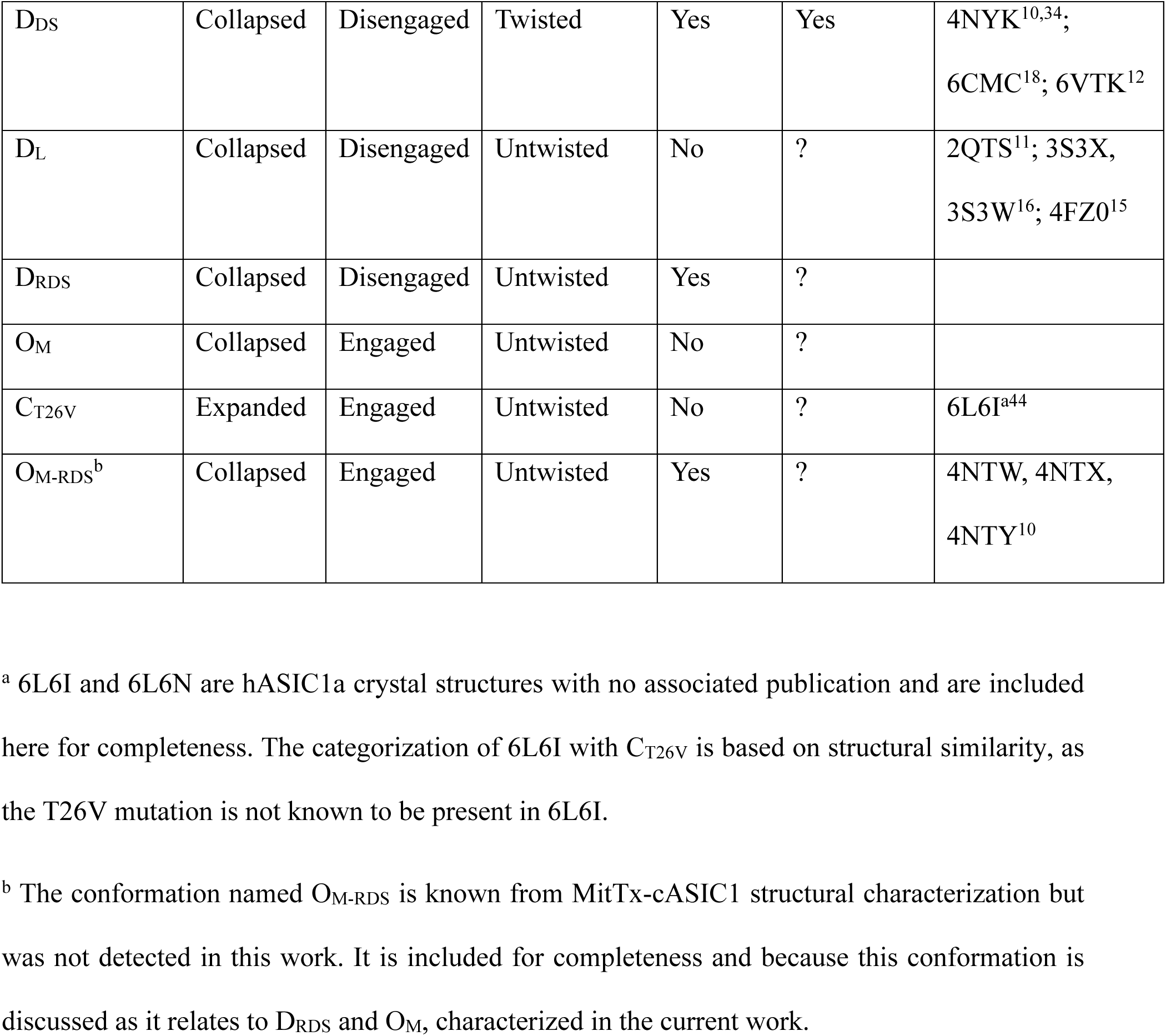
Summary of hASIC1a conformations.

| Conformation | Acidic pocket | Clutch | TMD arrangement | TM2b domain swap? | Re-entrant pre-TM1? | Similar to (PDB ID) |
| --- | --- | --- | --- | --- | --- | --- |
| C<sub>DS</sub> | Expanded | Engaged | Twisted | Yes | Yes | 5WKU, 5WKV, 6AVE<sup>13</sup>; 5WKX, 5WKY<sup>18</sup>; 6VTL<sup>12</sup>; 7CFS, 7CFT<sup>17</sup>; 6XH9<sup>20</sup>; 7RNN<sup>19</sup> |
| C<sub>A</sub> | Expanded | Engaged | Untwisted | ? | ? |  |
| O<sub>L</sub> | Collapsed | Engaged | Untwisted | No | ? | 4FZ1<sup>15</sup>; 6L6N<sup>a43</sup> |
| D<sub>DS</sub> | Collapsed | Disengaged | Twisted | Yes | Yes | 4NYK<sup>10,34</sup>;<br>6CMC<sup>18</sup>; 6VTK<sup>12</sup> |
| D<sub>L</sub> | Collapsed | Disengaged | Untwisted | No | ? | 2QTS<sup>11</sup>; 3S3X,<br>3S3W<sup>16</sup>; 4FZ0<sup>15</sup> |
| D<sub>RDS</sub> | Collapsed | Disengaged | Untwisted | Yes | ? |  |
| O<sub>M</sub> | Collapsed | Engaged | Untwisted | No | ? |  |
| C<sub>T26V</sub> | Expanded | Engaged | Untwisted | No | ? | 6L6I<sup>a44</sup> |
| O<sub>M-RDS</sub><sup>b</sup> | Collapsed | Engaged | Untwisted | Yes | ? | 4NTW, 4NTX,<br>4NTY<sup>10</sup> |

<sup>a</sup> 6L6I and 6L6N are hASIC1a crystal structures with no associated publication and are included here for completeness. The categorization of 6L6I with C<sub>T26V</sub> is based on structural similarity, as the T26V mutation is not known to be present in 6L6I.
<sup>b</sup> The conformation named O<sub>M-RDS</sub> is known from MitTx-cASIC1 structural characterization but was not detected in this work. It is included for completeness and because this conformation is discussed as it relates to D<sub>RDS</sub> and O<sub>M</sub>, characterized in the current work.

The first conformation, identified at pH 8.5 and 7.5, resembles the closed, resting state structures of cASIC1 and hASIC1a previously solved at high pH^12,17,19,20^. The three hASIC1a protomers exhibit threefold rotational symmetry around a central axis containing the putative pore. Each protomer’s ECD has a characteristic “hand holding a ball” architecture (Figure S3A). Within the ECD region, a relative expansion around the so-called acidic pocket has been attributed to deprotonation of acidic side chains creating an electrostatic repulsion^13^. The six TM helices form three parallel helical pairs; with the intracellular ends of each pair twisted in a clockwise direction around the pore axis. In this conformation, TM2 is not continuous; rather, it is domain-swapped, wherein the proximal half (TM2a) of one protomer abuts the distal half (TM2b) from another protomer with the GAS motif defining the separation distance of said elements. A 25-residue region directly N-terminal to TM1 is membrane-embedded in a “re-entrant loop” configuration (Figures S3B and S3C), positioning the highly conserved HG motif against the GAS motif in a hydrogen-bonding network proposed to confer structural stability to the domain-swapped conformation^12^. Overall, the resemblance to the previously resolved closed resting state structures and the domain-swapped architecture of TM2 led us to denote this conformation as C_DS_ (Table 1).

In addition to C_DS_, a second conformation was also observed at pH 7.5. The ECD conformation closely resembles C_DS_ (Figure S3A); however, the pairs of TM helices span the membrane without lateral displacement relative to the pore axis. This straightening can be characterized as an “untwisting” of the TMD. Because the GAS motif and TM2b cannot be resolved, it cannot be determined whether TM2 domain-swapping occurs in this conformation, and we have therefore categorized it as an “ambiguous” TMD conformation (C_A_). Although TM1 is resolved for a longer distance than TM2, pre-TM1 is not resolved and its conformation or position relative to TM1 cannot be determined.

A third major conformation was observed at both pH 7.5 and 6.5. Relative to both C_DS_ and C_A_, the ECD is in a more contracted configuration around the acidic pocket. This conformational change has been associated with protonation of the acidic side chains and neutralization of repulsive negative charges, and likely correlates with channel activation^11^. The TMD shares some features with C_A_, namely the untwisted TM helices relative to the pore axis; however, there are differences in the relative helical positionings. Additionally, the GAS motif and TM2b are resolved. This conformation has TM2a and TM2b forming a continuous, linearized TM2 helix, with the GAS motif wound into a single turn thereof. We therefore refer to this confirmation as O_L_, in line with functional evidence of open channels at these pHs (Figure S1). Pre-TM1 cannot be resolved, and the close positioning of the six TM helices does not leave a space for it to occupy a pore-lining “re-entrant loop” configuration as in C_DS_. The TMD additionally exhibits slight but noticeable asymmetry around the threefold axis of the ECD; as a result, O_L_ and all similar conformations lacking domain-swapped TM2bs (including C_A_) were analyzed using C1 symmetry instead of the C3 symmetry used for domain-swapped conformations such as C_DS_ (Table S3).

Three distinct conformations were observed at pH 5.7. The ECDs of these conformations resemble that of O_L_ in most respects. However, the characteristic inversion (“disengagement”) of the β11-β12 “clutch” linker, proposed to decouple the acid-sensing features of the ECD from the pore-forming TM2^13^, provides indication that the increased proton concentration has shifted the protein into a desensitized state. The three hASIC1a conformations obtained at pH 5.7 differ in TMD configuration. D_DS_ has a twisted, domain-swapped TMD configuration closely resembling the TMD in C_DS_, including the presence of pre-TM1 in its “re-entrant loop” configuration (Figure S3D). D_L_ has an untwisted TMD and linearized TM2, with a modest tilt of the TMD relative to the pseudo-threefold axis of the ECD that recalls the more dramatically tilted conformation of an early cASIC1 crystal structure solved at pH 5.5^11^. D_RDS_ has a unique TM configuration sharing some features of both D_DS_ and D_L_: it features an untwisted TMD and domain-swapped TM2, with outward splaying of TM2b and the lower part of TM1 relative to D_L_. Its name reflects that its TM helices are rotationally displaced relative to the common conformation of D_DS_ and C_DS_. The position of pre-TM1 in D_RDS_ is ambiguous; although there are weak map features beneath the GAS motif and between TM1 and TM2b consistent with the relative position of the “re-entrant loop” configuration of pre-TM1 in C_DS_ and D_DS_ (Figure S3E), their weakness in D_RDS_ precludes model-building or a confident assignment of this pre-TM1 configuration.

### MitTx-bound hASIC1a adopts a linearized transmembrane domain conformation

To provide further mechanistic context to the identification of O_L_ as the sole conformation observed at pH 6.5—a pH window putatively consistent with a relevant conducting state (Figure S1)—we resolved the structure of hASIC1a in complex with the snake venom toxin MitTx, a heteromeric toxin that activates ASIC1 channels independently of pH^7,10^. Conformation O_M_ resulted from the addition of MitTx to hASIC1a at pH 7.5 or 6.5 (Figures 2A and 2B; Figures S4A and S4B). We consider this conformation to be unique from O_L_ due to the influence of the extensive toxin-binding interface on the hASIC1a structure, but it bears strong overall resemblance to this conformation in both ECD and TMD (C_α_ positional RMSD for O_L-6.5_ vs. O_M-7.5_ 0.978 Å). This linearized TM2 configuration is unlike the domain-swapped arrangement observed in a crystal structure of MitTx-cASIC1^10^, which exhibits TM helical positionings similar to D_RDS_. The conformational discrepancy may relate to a species-specific phenomenon or a difference in experimental conditions. For example, whereas MitTx was added to cASIC1 at pH 7.4 in Tris^10,15^ (considered by the authors to be a non-activating pH condition), it was added to hASIC1a at either pH 7.5 or 6.5 in phosphate buffer, where a partial or full transition to C_L_ or O_L_ prior to MitTx addition might be expected based on structural analyses of hASIC1a without MitTx at these pH conditions (Figure 1).

**Figure 2:**
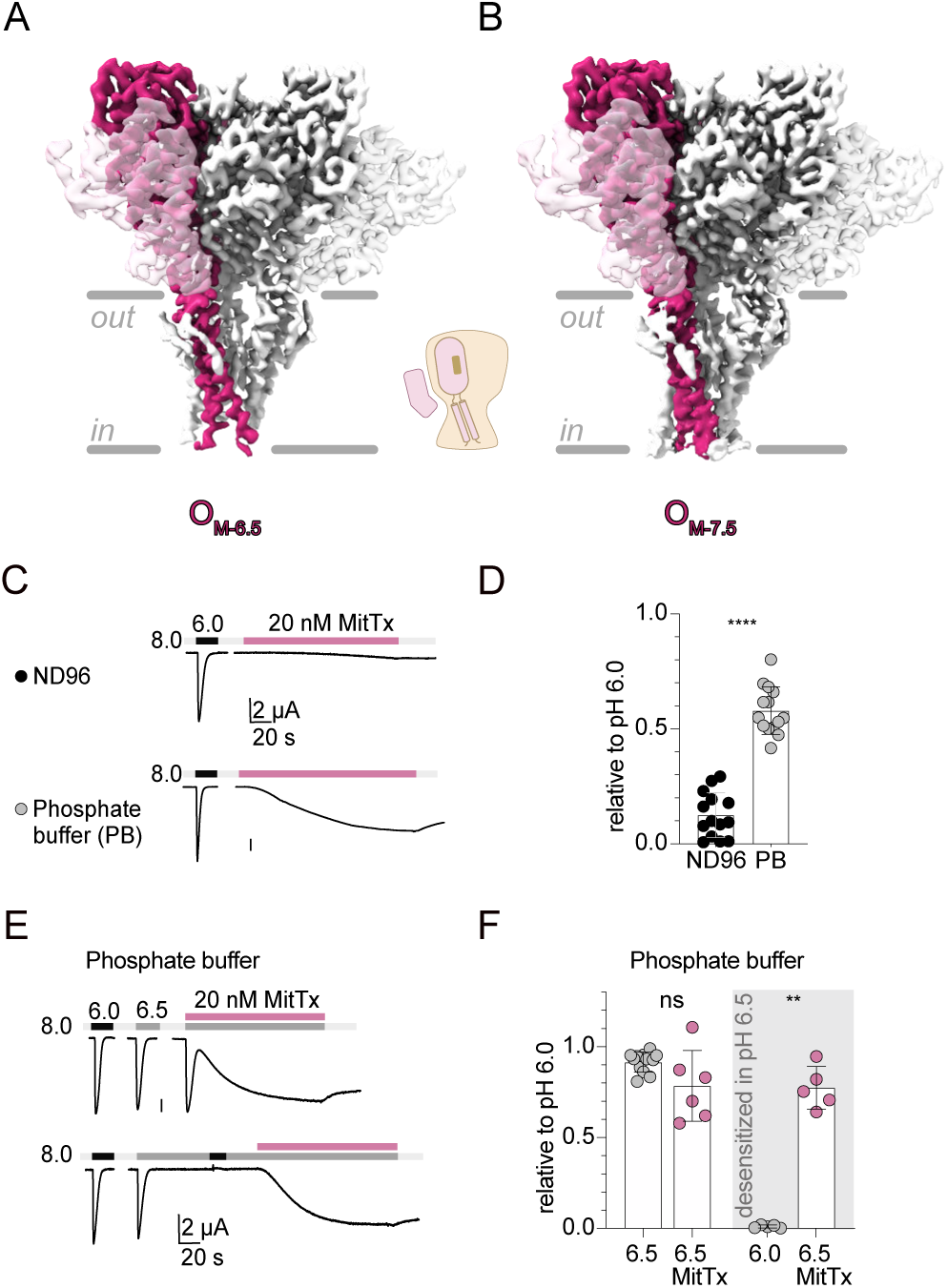
Modulation of hASIC1a by MitTx in phosphate buffer. **A-B**. Maps of O_M-6.5_ and O_M-7.5_. Map regions corresponding to an individual hASIC1a subunit are colored. Map regions corresponding to MitTx are shown in transparency. The orientation and approximate position of the homotrimer within the plasma membrane is indicated with gray lines. A cartoon schematic is shown between the two maps. **C.** TEVC traces showing the effect of 20 nM MitTx in ND96 (top) and phosphate buffer (bottom) at pH 8.0. Vertical scale bars are 2 μA; traces are on the same timescale. **D.** Normalized response of data shown in C comparing the response to 20 nm MitTx in ND96 and phosphate buffer (PB). **E.** TEVC traces showing the effect of 20 nM MitTx in phosphate buffer when applied at pH 6.5 (top) and when applied when channels were preconditioned at pH 6.5 before MitTx addition (bottom). **F.** Normalized responses of data shown in E. Responses to MitTx were reported at the plateau. Data are mean ± SD. Statistical analysis: non-parametric t-test *p<0.05, **p<0.005, ***p<0.0005, ****p<0.0001, ns=non-significant. See also Figure S4.

The functional characterization of the complex, comparing the effects of MitTx in either ND96 or phosphate buffer, revealed a greater activating effect of MitTx in phosphate buffer (Figures 2C and 2D; Table S1). When mimicking conditions under which purified hASIC1a was exposed to MitTx at pH 6.5, channels were first exposed to pH 6.5, prompting activation followed by desensitization. Subsequent addition of MitTx produced a sustained inward current, indicating that MitTx—unlike protons—can transition hASIC1a from a desensitized state to a stabilized conducting state (Figures 2E and 2F; Table S1).

The mechanism by which MitTx achieves this can be understood through its extensive interactions with hASIC1a. The mode of interaction has been characterized as a “church-key” that inserts bulky or charged side chains into structural features in the lower thumb, palm, and wrist domains to communicate outward leverage^10^. Each toxin interacts across the interface between two hASIC1a protomers, forming its most extensive (“major”) interaction with the hASIC1a subunit in the clockwise position (relative to the extracellular viewpoint) and a minor interaction with the counter-clockwise hASIC1a subunit. The side chain of MitTx-α-Phe14 packs into a pocket formed by the lower thumb domain of the major-interacting hASIC1a subunit and the β1-β2 linker of the minor-interacting hASIC1a subunit (Figure S4C); the β1-β2 linker itself interacts with the β11-β12 clutch that rearranges to decouple the ECD from the TMD in desensitized ASIC1^13^. The desensitization-associated “disengaged clutch” conformation of β11-β12 pulls β1-β2 away from the position which permits interaction with MitTx-α-Phe14, disrupting the pocket formed between hASIC1a subunits (Figure S4C). Considering the effects of MitTx binding on channel desensitization, the function of this specific MitTx-ASIC1a interaction may be to provide favorable contacts within the non-desensitized β1-β2/β11-β12 region and thereby energetically disfavor its rearrangement, while providing an energetic benefit to reverting clutch inversion when MitTx binds to desensitized protein.

### Mutation in the pre-TM1 region precludes TM2 domain-swapping at high pH

To further test the functional and conformational relevance of O_L_, we determined the structure of hASIC1a with a T26V mutation in the pre-TM1 region at pH 8.5 (Figure 3A; Figure S4D). Several mutations in the pre-TM1 region have been associated with perturbations in gating^21–23^ and divalent ion permeability^24^. This mutation was chosen because it had previously been shown to increase apparent proton sensitivity and display significantly slower desensitization kinetics in mouse ASIC1a^22^; similar results were observed with hASIC1a containing the T26V mutation (Figure 3B).

**Figure 3:**
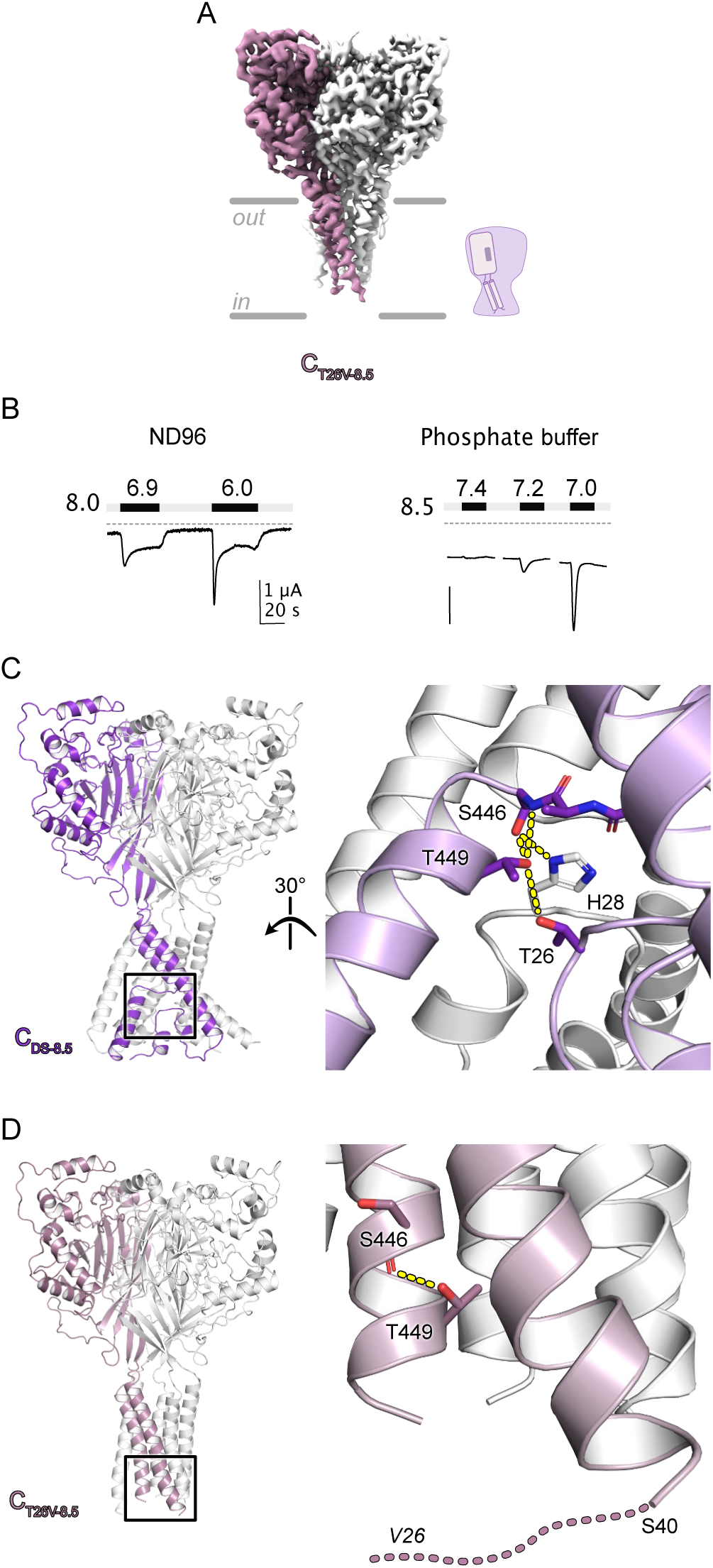
Functional and structural consequences of T26V mutation. **A**. Map of C_T26V-8.5_. Map regions corresponding to an individual hASIC1a subunit are colored. The orientation and approximate position of the homotrimer within the plasma membrane is indicated with gray lines. A cartoon schematic is shown beside the map. **B**. TEVC trace showing activation peaks of hASIC1a T26V in ND96 (left) and phosphate buffer (right). Vertical scale bars are 1 μA; traces are on the same timescale. The gray dashed line indicates 0 μΑ. In phosphate buffer, oocytes showed significant leak even at pH 8.5. **C**. The putative hydrogen-bonding network bridging Thr26, Thr449, Ser446 of the GAS motif, and His28 of the HG motif from a neighboring subunit is shown in the model of C_DS-8.5_. Side chains of involved residues and the full GAS motif are shown in stick representation. The putative H-bonding interaction network around Thr449 is indicated with dashed yellow lines. **D**. The position of Thr449 in the model of C_T26V-8.5_ is indicated in stick representation along with its putative H-bonding partner, the main-chain carbonyl oxygen of Ser446. Val26 is within the unmodeled N-terminal region, as represented with a dashed line extending from the first modeled residue (Ser40). See also Figure S4.

Introduction of the T26V mutation shifts the activation pH to more alkaline conditions in mouse ASIC1a^22^. We therefore reasoned it might shift the conformational equilibrium towards those typically observed at lower pH values. The single conformation observed at pH 8.5—named C_T26V_—most closely resembles C_A_; however, due to higher map quality within the TMD, TM2 can be stated unambiguously to be in the linearized conformation. It is not possible to determine whether C_T26V_ and C_A_ represent the same conformation due to the importance of TM2b to this interpretation and the relatively modest overall resolution of C_A_; however, the ECD, TM1, and TM2a of C_T26V_ closely resemble those of C_A_ (C_α_ positional RMSD for C_A-7.5_ vs. C_T26V_ 0.660 Å). Their close similarity suggests that the T26V mutation stabilizes the C_A_ conformation that is accessed by wild-type protein.

The loss of the “re-entrant loop” configuration of pre-TM1 at pH 8.5 appears to be a direct consequence of the single isosteric T26V mutation, which removes hydrogen-bonding capability from the formerly polar side chain. When wild-type protein is in the C_DS_ conformation, the Thr26 side chain forms a hydrogen bond with the side chain of Thr449, and the T26S^22^ (but not the T26C)^22,23^ mutant retains wild-type-like function. Thus, the loss of this single hydrogen-bonding interaction may be sufficiently destabilizing to the “re-entrant loop” configuration of pre-TM1 to induce a full conformational shift to C_T26V_.

In C_DS_, Thr449 and Thr26 participate in a larger hydrogen-bonding network that connects two highly conserved elements: Ser446 of the GAS motif and His28 of the HG motif (the latter originating from a neighboring protomer) (Figure 3C). Therefore, conformational destabilization of the Thr449 hydroxyl by removal of the Thr26 hydroxyl may communicate this destabilization into functionally crucial residues of hASIC1a with well-characterized structural roles in C_DS_. In C_T26V_, the Thr449 side chain cannot maintain any of its C_DS_-specific hydrogen-bonding interactions, although the side chain hydroxyl retains proximity to Ser446 via the main-chain carbonyl of the latter (Figure 3D). Because the conformation of pre-TM1 in C_T26V_ is not known, nor can it be observed in any of the other hASIC1a conformations with continuous TM2s, it is possible that there are further structural consequences of the T26V mutation beyond the observed conformational shift at pH 8.5. For example, Thr26 in C_A_, O_L_, or D_L_ may engage in other specific interactions that are also disrupted by the T26V mutation, contributing to the observed functional perturbations.

In functional experiments in phosphate buffer, *Xenopus* oocytes expressing the T26V mutant exhibited significant leak, which appeared largely pH-independent between pH 8.5 and 7.4 (Figure 3B). Although activation was observed at pH 7.2, reliable characterization of activation was hindered by the noticeable leak and pronounced tachyphylaxis in some oocytes. However, the phenotypic perturbations suggest that C_T26V_ readily transitions into an open state, a condition possibly exacerbated by phosphate buffer.

### Modeled pore characteristics support assignments of functional states

Although well-documented structural characteristics of the ECD and functional analyses were used to categorize the structures according to their putative functional states, we further analyzed the molecular models to interpret the likely ability of sodium ions to access and travel through the pore. Fenestrations between the ECD-proximal ends of each TM1 and its neighboring subunit’s TM2a are a proposed access site due to the lack of evidence for release of constriction points along the threefold axis of the ECD. These windows extend from the “wrist” region between ECD and TMD deep into the membrane-embedded portion of the protein, with evidence of lipid-like features occupying the spaces within the transmembrane region (Figure 4A). The widest fenestrations are found in O_L_, D_RDS_, and O_M_. They are slightly asymmetrically narrowed in D_L_, and sharply narrowed in C_DS_, C_A_, D_DS_, and C_T26V_ (Figures S5A-S5H). Restricted access to the pore due to narrow fenestrations is consistent with nonconducting channels, although conformations with wide fenestrations cannot be concluded to be conducting states due to the possibility of constrictions within the pore itself.

**Figure 4:**
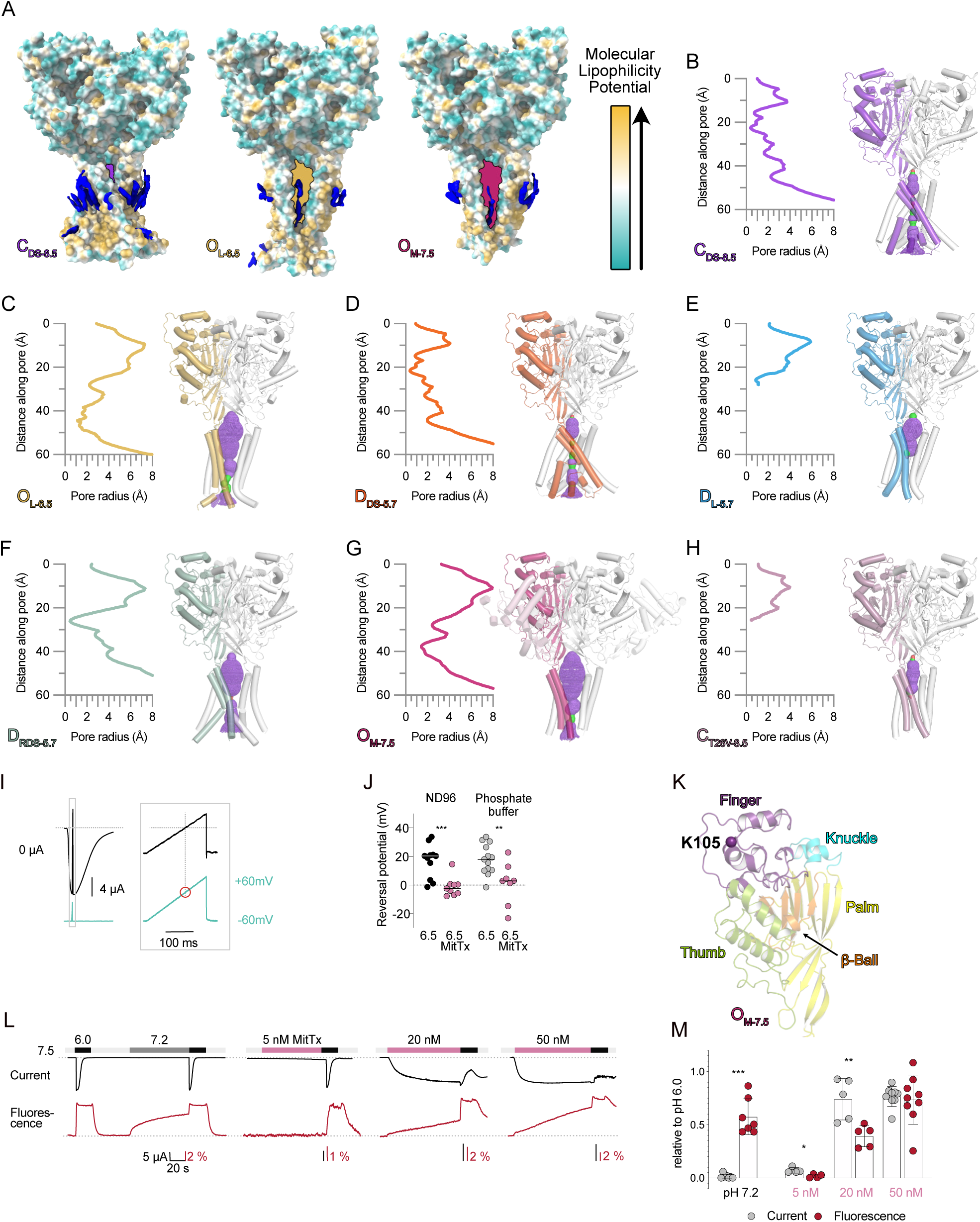
Pore accessibility of hASIC1a conformations. **A**. Models of C_DS-8.5_ (left), O_L-6.5_ (center), and O_M-7.5_ (right) are depicted in surface view colored by molecular lipophilicity potential. Map features in the TMD region that appear to correspond to neither micelle nor protein are shown in blue. Spaces between TM1 and TM2 of adjacent subunits (fenestrations) are drawn in colored shapes. For O_M-7.5_, surface corresponding to MitTx is omitted for clarity. **B-H**. Pore profile analyses of the indicated model were performed using HOLE. Pore radius plots (left) and pore maps (right) are shown starting from the consensus C_α_ position of Val74 at the base of the palm domain. For calculation of pore profiles depicted in E and H, the starting point was set to the centroidal position of Leu441 to approximate the trajectory of an ion that traveled down the pore, encountered the impassable constriction point, and stopped; otherwise the starting point was set to coordinates at the approximate extracellular plasma membrane interface. Pore maps are colored according to radius: red < 1.15 Å < green < 2.3 Å < purple. **I**. TEVC trace showing how the reversal potential was established by running a voltage-ramp protocol from -60 to +60 mV within 200 ms near the peak of a pH 6.5 activation. **J.** Comparison of reversal potentials when channels were activated using pH 6.5, or at the plateau of the response to 20 nM MitTx in pH 6.5 in ND96 (left) or phosphate buffer (right). **K**. Structure showing position Lys105, where an introduced cysteine was used for fluorescent labeling. **L.** Voltage-clamp fluorometry traces of hASIC1a K105C, labeled with Alexa Fluor 488 via a maleimide linker, with currents in black and fluorescence in red. Channels were activated with pH 6.0, conditioned with pH 7.2, and exposed to different concentrations of MitTx at pH 7.5 in phosphate buffer. All traces are on the same timescale and vertical scale bars are 5 μA. **M.** Normalized responses from L. Data are mean ± SD. Statistical analysis: non-parametric t-test *p<0.05, **p<0.005, ***p<0.0005, ns=non-significant.

Further interpretation of structural determinants of channel conductance was done through pore profile analyses using HOLE^25^ (Figures 4B-4H; Figures S5I-S5K). This analysis was limited in some cases by the lack of visible side chain features. Although the helical turns were generally clear and the side chain progression along a regular α-helix could be used as a guide to line the pore appropriately, the specific orientations of many pore-lining side chains could not be determined. However, the general trends in pore profile can be determined with reasonable confidence.

The pore is lined entirely by TM2 residues, except when pre-TM1 is in its “re-entrant loop” configuration and its residues line the lower ion permeation pathway. C_DS_ and D_DS_ have constrictions around Asp434 and Gly437—despite the contribution of pre-TM1 to the pore in these conformations, the tightest constrictions are associated with residues of TM2a (Figures 4B and 4D; Figure S5I). Combined with the narrow fenestrations, the pore features are generally consistent with the assignments of their cASIC1 equivalents as, respectively, a high-pH closed resting state^12,17,19,20^ and a nonconducting desensitized state^12,18^.

In other conformations, Leu441 appears to be the major site of constriction. Despite relatively wide fenestrations, D_L_ and D_RDS_ have very tight constrictions around Leu441 (Figures 4E and 4F); C_T26V_ combines narrow fenestrations and a tight constriction around Leu441 (Figure 4H). Due to insufficient map quality within the TM region, C_A_ was omitted from pore profile analysis. However, the overall agreement in helical alignment of TM1 and TM2a between C_A_ and C_T26V_ presents a likelihood that C_A_ is similarly constricted around Leu441. This position has previously been identified as an important site for normal function of ASIC1; alanine substitutions at this position are associated with gating perturbations including reduced ion selectivity^26,27^.

O_L_ and O_M_ are therefore left as the only conformations with wide fenestrations and relatively unconstricted pores (Figures 4C and 4G; Figures S5J and S5K). Although both exhibit degrees of pore narrowing in the TM2b region, the constrictions are less dramatic than in other conformations. These constricted areas are asymmetric relative to the pseudo-threefold axis (especially in O_L_) and variable between their representative maps. We consider the possibility that the distal TMD in O_L_ and O_M_—and, for similar reasons, in the other conformations with continuous TM2s—is inherently more flexible than in the conformations with domain swapped TM2s and “re-entrant loop” pre-TM1. This flexibility may also increase the susceptibility of this region to distortions due to the biochemical environment of the detergent-extracted protein, potentially inducing an apparent narrowing of the pore.

Although the O_M_ pore profile is only slightly different from that of O_L_ (Figures 4C and 4G; Figures S5J and S5K), functional experiments with MitTx-activated hASIC1a suggest a decreased sodium ion selectivity compared to proton-activated hASIC1a (Figures 4I and 4J). In comparison, cASIC1 was shown to retain some ion selectivity with MitTx-induced activation, although to a lesser degree than proton-activated cASIC1^10^. Lost ion selectivity relative to proton-activated hASIC1a suggests perturbation of the selectivity filter, but the overall similarity of O_L_ and O_M_ indicates that the perturbation may be subtle or rely on details that are not adequately represented in these models. Because local resolution limitations in the TMD region (Figures S2D and S2E; Figures S4A and S4B) resulted in ambiguity in the orientational positioning of many pore-lining side chains, the precise structural differences that contribute to this functional perturbation may have become obscured.

The small structural perturbations of O_L_ *versus* O_M_ and the ability of MitTx to induce a conducting state from either the resting or desensitized state of hASIC1a support the argument that MitTx-induced activation is distinct from activation elicited by protons. To test this notion directly, we turned to voltage-clamp fluorometry (VCF), which allowed us to monitor conformational rearrangements around the Lys105 reporter site in response to activation by MitTx or protons^28–30^ (Figure 4K). While even subactivating pH conditions lead to changes in fluorescence in this position and proton activation results in rapid and maximal change in fluorescence, activation of the channels by MitTx leads to smaller and slower-developing changes in fluorescence (Figures 4L and 4M; Table S1). Together, these data indicate that the mechanism of activation by MitTx is indeed different to that of protons, even if the final open state bears strong resemblance.

Considering the proposed increase in TMD flexibility associated with O_L_ and O_M_, we suggest that the T26V mutation similarly increases the flexibility of the TM helices at high pH due to the destabilization of the “re-entrant loop” configuration of pre-TM1. Although the constricted pore profile and narrow fenestration shape of the C_T26V_ model (Figure 4H; Figure S5D) are not consistent with C_T26V_ itself representing an open state, increased TMD flexibility relative to C_DS_ would likely make the mutant protein more prone to adopt an open conducting state. The openings at high pH may be transient in nature, and therefore not identified by structural characterization, but may nevertheless explain the leak currents observed with T26V hASIC1a in phosphate buffer. This interpretation would suggest that C_T26V_ represents a destabilized closed state at pH 8.5, resembling a pre-open state with a low barrier to transitioning into a conducting state.

### Unexpected conformational diversity in desensitized states

ASIC1a function normally involves rapid and complete desensitization after proton-induced activation^31^. A particularly dramatic structural feature of desensitization involves a conformational rearrangement of the β11-β12 linker with an inversion of Leu415 and Asn416 and concomitant rearrangement of the neighboring β1-β2 linker (Figures 5A-5C; Figure S6). Functional experiments have also highlighted the importance of the β11-β12 and β1-β2 linkers for desensitization32^,33^. In conformations observed at pH 5.7, all ECDs present the characteristic collapsed acidic pocket and disengaged clutch associated with the desensitized state, but the TMs feature remarkable conformational diversity (Figures 5A and 5D).

**Figure 5:**
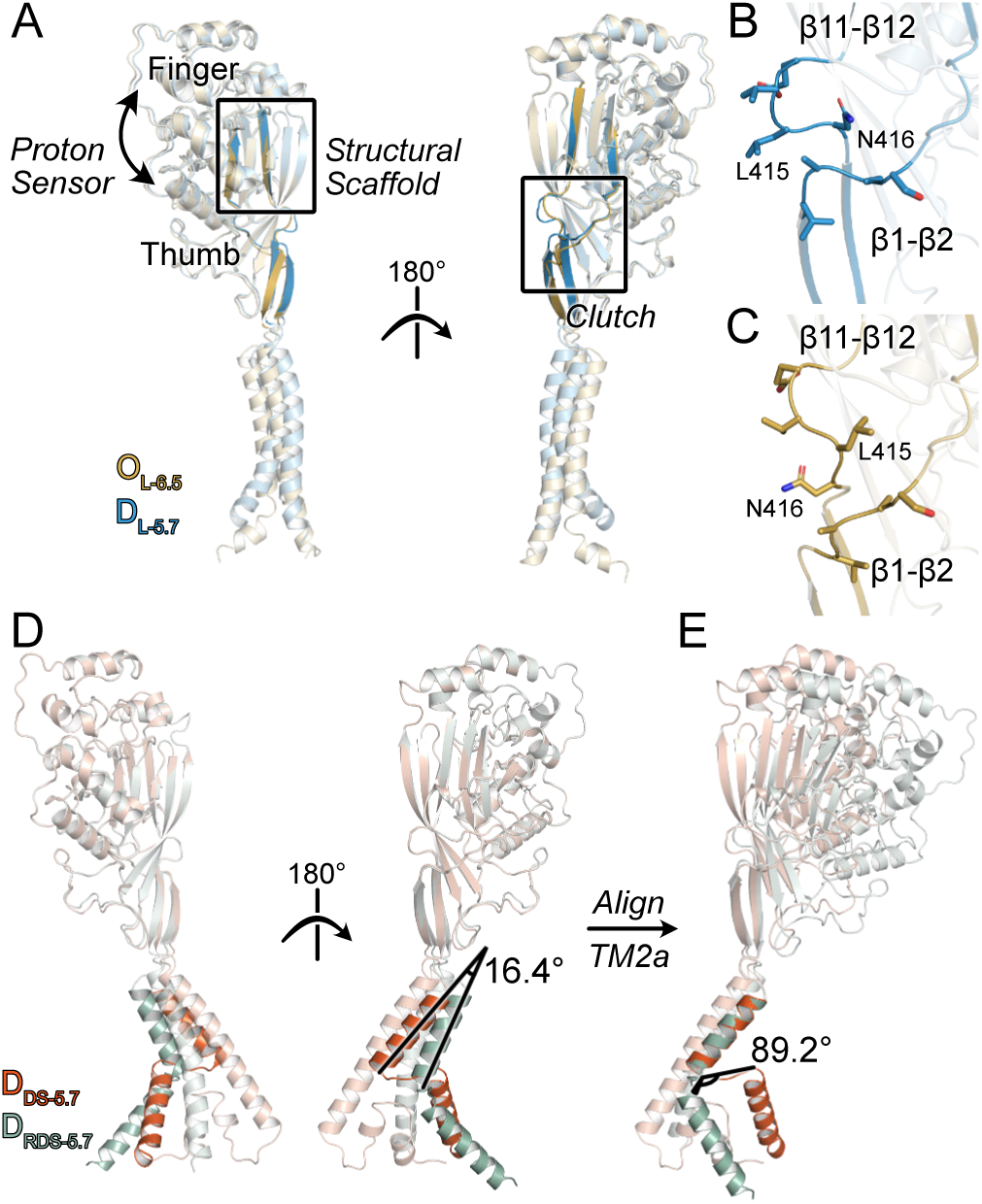
Three desensitized conformations of hASIC1a. **A.** Superposition of a single subunit from D_L-5.7_ and O_L-6.5_. ECD structural elements are indicated over models. The desensitization-associated “clutch” (right panel) is highlighted in relation to the structural scaffold of the upper palm domain and the proton-responsive thumb and finger domains. **B-C**. Close-up views of the clutch region of, respectively, D_L-5.7_ (B) and O_L-6.5_ (C). **D**. Relative to D_DS-5.7_, the rotational displacement of TM2a from D_RDS-5.7_ is noted. **E**. TM2a of D_RDS-5.7_ was aligned to TM2a of D_DS-5.7_ to isolate and measure the rotational displacement of the GAS motif. See also Figure S6.

While it is unclear in isolation whether ASIC1a transitions directly between D_RDS_ and either D_DS_ or D_L_—and, if so, in which order they are accessed—the rotational differences between D_RDS_ and D_DS_ are best represented by describing the transitions necessary to move between them. The conformational rearrangement required to transition from D_RDS_ to D_DS_ involves a clockwise twisting (relative to the intracellular view) of both TM1 and TM2. The rotational displacement of TM2 between D_RDS_ and D_DS_ involves a 16.4° upward swing of TM2a and an 89.2° rotation of the GAS motif and TM2b within the plane of the membrane (Figures 5D and 5E).

Structural characterizations of cASIC1 in the “disengaged clutch” conformation have previously described either domain-swapped^10,12,18,34^ or continuous^11,15,16^ TM2s; D_DS_ bears resemblance to the former and D_L_ to the latter in their respective TMD regions. Although there is no direct cASIC1 analogue to D_RDS_ in the library of solved structures, the crystal structure of MitTx-bound cASIC1^10^ has strong similarities in the TMD region; specifically in the otherwise unique TM rotational displacements of D_RDS_ relative to D_DS_. However, MitTx-cASIC1 has the ECD structural markers of a non-desensitized state and slightly different TM helical positions, including a lateral displacement of TM2a that avoids the major pore constriction predicted in D_RDS_ due to displacement of Leu440 (the cASIC1 equivalent of Leu441) away from the pore axis. Together, these findings suggest that desensitization of hASIC1a involves multiple nonconducting TMD conformations, underscoring the striking degree of TMD conformational variability observed across the functional states of hASIC1a.

## Discussion

In this study, we have used complementary structural, functional and fluorometric approaches to establish that hASIC1a open conformations are associated with linear TM helix conformations, while the TMD can adopt a surprisingly diverse array of conformations in desensitized states. We further provide direct evidence for mechanistic differences between proton and toxin activation, as well as for how pre-TM1 forms interactions that stabilize the closed channel. Together, these results establish a firm framework to understand the structural underpinnings of ASIC gating, which have long remained elusive.

### Implications for ASIC gating

Functionally speaking, ASIC1 has at least three discrete states: a resting state, an activated state, and a desensitized state. However, there is evidence that the actual landscape of functionally relevant structural states is more complex than three conformations could explain^35–37^. In general, assigning structures to functional states has been inferential, based on the agreement of conditions used for electrophysiological, biophysical, and structural studies such as pH modulation and toxin addition. By varying pH between 5.7 and 8.5, we have captured six different conformations with four distinct TMD architectures: two with a twisted TMD, domain-swapped TM2bs, and “re-entrant loop” pre-TM1s in resting and desensitized conformations (C_DS_ and D_DS_); two with continuous TM2bs in an untwisted arrangement, in conducting and desensitized conformations (O_L_ and D_L_); one desensitized conformation with untwisted TMD and domain-swapped TM2bs (D_RDS_); and one high-pH conformation with untwisted TMD but unknown TM2b architecture (C_A_). Additional conformations resembling, respectively, C_A_ and O_L_ were captured through probing the modulatory effects of a gating-related mutation (C_T26V_) and agonist toxin MitTx (O_M_); based on functional characterizations, the T26V mutation and MitTx binding have non-identical functional consequences, but both are associated with increased pH sensitivity, decreased ion selectivity^22^, and reduced desensitization rates relative to wild-type protein without MitTx. Taking together the eleven structural models representing six major conformations of hASIC1a, we propose an overall mechanistic model of conformational rearrangements upon activation (pH- or toxin-induced) and desensitization (Figure 6).

**Figure 6:**
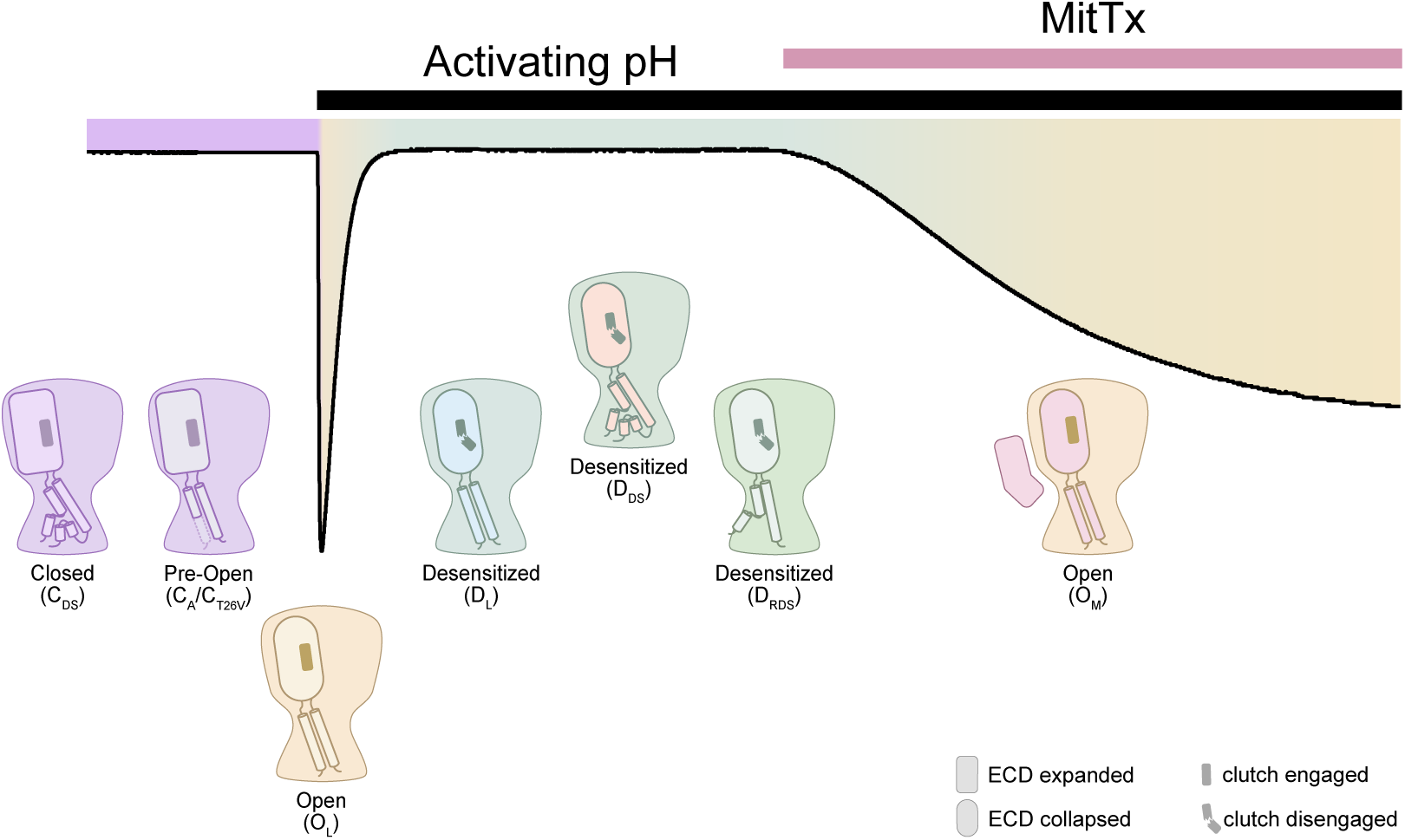
Proposed distribution of distinct hASIC1a conformations along a hASIC1a current trace. At high pH, hASIC1a adopts a non-conducting, resting conformation (C_DS_), and a closed pre-open state (C_A_/C_T26V_) with a lower transition barrier to the open state. The channel opens at low pH (O_L_) and rapidly desensitizes into several states (D_RDS_, D_DS_, and D_L_). MitTx can activate hASIC1a (O_M_) from the desensitized (shown) or closed states, through a mechanism distinct from pH-dependent activation.

As domain-swapped TMD conformations were observed at both high and low pH but neither at pH 6.5 nor in the presence of MitTx, adopting a linearized-TM2 conformation is likely a conformational bottleneck associated with the transition into a conducting state. In this model, the three discrete desensitized states observed could be interpreted as sequential stages in the relaxation toward the domain-swapped TMD conformation associated with the closed resting state. The series of events would then involve the following steps: the channel starts in the closed resting state with domain-swapped TMD (C_DS_), from which mild acidification may shift the TMD conformation toward another non-conducting state possessing characteristics of a pre-open state (C_A_). From there, acidification induces ECD perturbations, such as collapse of the acidic pocket due to acidic residue protonation, along with full TMD rearrangement into a linearized conformation (O_L_). Prolonged acidification then leads to inversion of the β1-β2 and β11-β12 linkers into the desensitization-associated “disengaged clutch” ECD conformation with minor inward shifting of TM1 and TM2 (D_L_), followed by unwinding of the GAS motif and reformation of the TM2 domain-swap with minor outward displacement of TM1 to accommodate the movement of TM2b (D_RDS_), and movement of the TMD back into the twisted, domain-swapped conformation (D_DS_). This state would then be poised to undo the protonation-associated ECD shifts and return to the closed resting state conformation (C_DS_) upon neutralization of the acidified extracellular environment.

As an alternative to a model in which each structurally captured desensitized state is visited in sequence, each of the three states might instead be visited directly from the resting or open state. The conformational diversity of the hASIC1a TMD at pH 5.7 recalls functional experiments that similarly are best explained by multiple species of desensitized ASIC1a. These functionally distinct desensitized states include an acute and rapidly reversible one, which may itself be divided into multiple subpopulations to explain the observation of multiple kinetic phases of acute desensitization in ASICs^9^ (Figure S1E). Another characterized desensitization state is long-lived, accumulates through repeated stimulation, and results in an incremental reduction in response (tachyphylaxis)^35–38^. An additional desensitization mechanism has been proposed that bypasses activation and is induced directly from the closed resting state by prolonged incubation at pHs slightly higher than the activation window (SSD)^39^. The coexistence of multiple structural states in the same ensemble presents a possibility that one or more of these conformations is the unique structural manifestation of one of these mechanistically relevant desensitization mechanisms, although the methods used here do not provide direct evidence in support of such assignments. As the most complete D_DS_-like cASIC1 structure was solved at a putative sub-activating pH condition (pH 7), it was proposed to reflect the conformation associated with SSD^12^; its identification as one of several hASIC1a conformations (D_DS_) induced by incubation at pH 5.7 may facilitate an increased understanding of the nature of the cASIC1 conformation or represent an alternative pathway to an equivalent structural state.

### Potential Methodological Constraints

Interestingly, conformations resembling desensitized states were observed only at pH 5.7. While the activation curve suggests that the majority of channels will access the open state at pH 6.5 (Figures S1B and S1F), the recorded current indicates that rapid and complete desensitization occurs. Therefore, one might expect an accumulation of desensitized protein within the samples used for grid preparation, which was not observed. This discrepancy may arise from differences between the buffers used for recording and grid preparation or be a consequence of detergent extraction of hASIC1a, potentially stabilizing a particular conformation that is less persistent in the environment of the cell membrane.

We recognize that the use of digitonin may influence hASIC1a’s conformational transitions in response to varying proton concentrations, and that phosphate buffering systems have functional consequences such as the observed shifts in the activation and SSD curves (Figure S1). However, this digitonin-solubilized protein has reproduced equivalents of cASIC1a structures solved in a membrane environment stabilized by styrene-maleic acid copolymer^12^ at three of the four sampled pH conditions (C_DS-8.5_, C_DS-7.5_, and D_DS-5.7_), supporting the use of digitonin as an effective reagent for capturing the range of hASIC1a conformations. Moreover, our structural examination of a single hASIC1a construct using a consistent buffering system captured most of the conformations previously seen in cASIC1 and others not previously observed in cASIC1 (Table 1), demonstrating both the versatility of the method when interrogating the hASIC1a conformational space and the inherent conformational variability of the TMDs. Future experiments employing technologies that preserve a more native membrane environment might further elucidate ASIC gating mechanisms.

### Outlook

The six major conformations captured in this study provide detailed mechanistic insights into ASIC1a function, highlighting an unexpected intrinsic conformational variability of the TMD. These conformational rearrangements are mediated by proton or toxin binding to the ECD to produce functionally distinct states, albeit suggesting linear TMD helices as a common denominator for otherwise slightly divergent open states. By comparison, less is known about structural transitions and conformational plasticity in other members of the broader ENaC/degenerin family, but recent work suggests that there might be important differences in, e.g., ENaCs^22,40^ and neuropeptide-activated FaNaCs (FMRFamide-activated sodium channels)^41,42^.

## Supporting information

Key resources table

## Acknowledgements

We thank members of the Baconguis and Gouaux labs for discussion on this study. A portion of this research was supported by NIH grant R24GM154185 and performed at the Pacific Northwest Center for Cryo-EM (PNCC) with assistance from Marcelo De Farias, Theo Humphreys, Janette Myers, and Rose Marie Yasuda. We thank Rui Yan at the HHMI Janelia CryoEM Facility for help in microscope operation and data collection. All structural experiments included in this study were supported by a grant from the NIH to I.B. (GM138862). Work in the Pless group was supported by Lundbeck Foundation grant R313-2019-571.

## Author Contributions

J.C., S.A.H., M.H.P., S.A.P., and I.B. designed experiments; J.C. did protein expression, purification, and grid preparation for wild-type hASIC1a; K.A.H. did protein expression, purification, and grid preparation for mutant (T26V) hASIC1a; S.A.H. did electrophysiological (TEVC) and voltage clamp fluorometry experiments; M.H.P. did electrophysiological (whole-cell patch clamp) experiments; J.C., K.A.H., and C.Y. processed cryo-EM data; K.A.H. built and analyzed structural models; J.C. wrote the initial draft manuscript; K.A.H. wrote the manuscript with contributions from J.C., S.A.H., M.H.P., S.A.P., and I.B.; S.A.P. and I.B. supervised research and acquired funding.

## Supplemental information titles and legends

Document S1. Figures S1-S7 and Tables S1-S4.

## Supplementary Figure Legends

**Figure S1: Functional characterization of hASIC1a in phosphate buffer by electrophysiology, related to Figure 1**

**A.** Representative excerpts of two-electrode voltage clamp (TEVC) traces of oocytes expressing hASIC1a activated by decreasing pHs of ND96 (top) or phosphate buffer (bottom). **B**. Normalized activation responses in ND96 and phosphate buffer. **C**. TEVC traces showing the induction of steady-state desensitization by preconditioning the channel for 2 min in decreasing pHs of ND96 (top) or phosphate buffer (bottom) before activation. **D**. Normalized SSD responses in ND96 and phosphate buffer. **E**. Representative trace from HEK293T* cells expressing hASIC1a, recorded using fast perfusion patch clamp electrophysiology in phosphate buffer. The hASIC1a current in phosphate buffer shows the typical fast activation and desensitization at pH 6 and an atypical standing current at pH 7.5. **F**. Normalized activation response of hASIC1a WT expressed in HEK293T* cells, recorded in whole-cell patch-clamp experiments. **G**. Graph summarizing the desensitization rates for hASIC1a in regular physiological solution (black circles) and in phosphate buffer (grey circles). In A and C, vertical scale bars are 2 μA, all traces are on the same timescale. Data in B, D, and F are mean ± SD.

**Figure S2: Cryo-EM data processing of hASIC1a maps attained with pH modulation, related to Figure 1**

**A-H**. Local resolution estimation (top), orientation distribution (middle), and Fourier shell correlation (FSC) (bottom) for each indicated map. The resolution at the FSC=0.143 cutoff (black line) is indicated.

**Figure S3: Structural overview of hASIC1a, related to Figure 1**

**A.** Cartoon representation of (left) C_DS-8.5_ and (right) O_L-6.5_ with color-coding according to the “hand holding a ball” model of subdomain architecture. **B-E**. Map regions corresponding to TM1/pre-TM1 (blue) and TM2 (orange) are shown in mesh representation over the model in stick representation. For E, unmodeled map features in the approximate expected position of the pre- TM1 “re-entrant loop” are shown.

**Figure S4: Cryo-EM data processing of hASIC1a maps attained using toxin addition or mutagenesis, related to Figures 2 and 3**

**A-B**. Local resolution estimation (top), orientation distribution (middle), and Fourier shell correlation (bottom) for the maps of hASIC1a in complex with MitTx at pH 6.5 and 7.5. The resolution at the FSC=0.143 cutoff (black line) is indicated. **C**. Left panel: Reference image of O_M-7.5_ with a highlighted region corresponding to panels on the right. Middle panel: The positioning of MitTx-α-Phe14 in the model of O_M-7.5_ is shown relative to the β1-β2 linkers of the minor-interacting hASIC1a in the models of O_M-7.5_ (magenta) and D_L-5.7_ (blue). Right panel top: The packing of the MitTx-α-Phe14 side chain (pink) between the β1-β2 linker of the minor-interacting hASIC1a subunit (magenta) and the α5-β10 loop of the major-interacting hASIC1a subunit (white) is represented with surface over a cartoon representation. Arrows indicate the proximity of the two hASIC1a subunits within the depicted region. Right panel bottom: The position of the MitTx-α-Phe14 side chain from the aligned O_M-7.5_ model is shown in the context of the D_L-5.7_ model. Arrows indicate the relative reduction in proximity between the β1-β2 linker and α5-β10 loop in D_L-5.7_. Inset in each panel: The spatial relationship between the colored hASIC1a and MitTx subunits is represented using a simplified surface view from the extracellular viewpoint. **D**. Local resolution estimation (left), orientation distribution (middle), and Fourier shell correlation (right) for the map of hASIC1a-T26V at pH 8.5. The resolution at the FSC=0.143 cutoff (black line) is indicated.

**Figure S5: Fenestrations and pore profiles of hASIC1a models, related to Figure 4**

**A-E**. Indicated models are depicted in surface view colored by molecular lipophilicity potential. Spaces between TM1 and TM2 of adjacent subunits (fenestrations) are drawn in colored shapes. For O_M-7.5_, surface corresponding to MitTx is omitted for clarity. Three rotational views are shown for each model. **F-H**. Models and fenestrations are depicted as in A-E. As the maps for these models were calculated with C3 symmetry imposed, only a single orientation is depicted. **I-K** Pore profile analyses of the indicated model were performed using HOLE. Pore radius plots (left) and pore maps (right) are shown from the consensus C_α_ position of Val74 at the base of the palm domain. For calculation of pore profiles, the starting point was set to coordinates at the approximate extracellular plasma membrane interface. Pore maps are colored according to radius: red < 1.15 Å < green < 2.3 Å < purple.

**Figure S6: Map quality of hASIC1a in the β1/β2 and β11/β12 regions, related to Figure 5**

**A.** The positions of the β1, β2, β11, and β12 sheets of D_L-5.7_ are shown in stick representation with map mesh overlay. Proximal structural motifs are labeled. **B.** The β1-β2 and β11-β12 sheets and linkers of D_L-5.7_ are shown in stick representation with map mesh overlay. **C**. The β1-β2 and β11-β12 sheets and linkers of O_L-6.5_ are shown in stick representation with map mesh overlay.

**Figure S7: Sample preparation and cryo-EM data processing strategy, related to STAR Methods**

**A.** The hASIC1a expression construct has a N-terminal 8xHis-eGFP tag which can be removed by thrombin cleavage. **B**. FSEC chromatogram of proteins extracted from HEK293S GnTI^-^ cells expressing GFP-tagged hASIC1a. GFP fluorescence is monitored due to the abundance of non-fluorescent cellular proteins within the sample. **C**. FSEC chromatogram of purified hASIC1a demonstrates a homogenous sample. Trp fluorescence is monitored due to removal of GFP during purification. **D**. SDS-PAGE gel of purified hASIC1a. The larger band at approximately 65 kDa is consistent with the size of hASIC1a; the bands at approximately 50 and 15 kDa were identified as hASIC1a fragments by N-terminal sequencing. **E-F**. Maps of C_DS-8.5_ and O_M-7.5_ (blue mesh) show weakness in the vicinity of Ser146, which is the N-terminal residue of the post-cleavage fragment. However, maps depict the side chains of neighboring Phe144 and Phe147, anchoring the model such that an intact Phe-Arg-Ser-Phe peptide can be modeled in spite of potential partial cleavage. Inset: The cleavage site vicinity is depicted as sticks in the context of the hASIC1a trimer. **G.** Although no cleavage was suggested to occur between Arg146 and Asn147 of cASIC1^12^, a similar map weakness is shown in desensitized cASIC1 (PDB ID: 6VTK) despite otherwise high map quality in the region. **H**. A workflow schematic for cryo-EM data processing using Cryosparc. The indicated data processing strategy resulted in maps for C_DS-7.5_, C_A-7.5_, and O_L-7.5_.

## Supplementary Table legends

**Table S1. Summary of electrophysiology data, related to Figures 2, 4, and S1**

**Table S2. PDB and EMDB codes of hASIC1a structures, related to Figures 1, 2, and 3**

**Table S3. Cryo-EM data collection, refinement, and validation statistics, related to Figures 1, 2, and 3**

**Table S4. Model refinement and validation statistics, related to Figures 1, 2, and 3**

## STAR Methods

### Resource availability

#### Lead contact

Further information and requests for resources and reagents should be directed to and will be fulfilled by the lead contact, Isabelle Baconguis.

### Materials availability

For any correspondence or requests related to the materials used in this study, please contact the lead contact.

### Data and code availability

- Cryo-EM maps and atomic models have been deposited in the Protein Data Bank and Electron Microscopy Data Bank and are publicly available as of the date of publication. The PDB IDs are 9E4A, 9E4B, 9E4C, 9E4D, 9E4E, 9E4F, 9E4G, 9E4H, 9E4I, 9E4J, and 9E4K. The EMDB IDs are EMD-47503, EMD-47504, EMD-47505, EMD-47506, EMD-47507, EMD-47508, EMD-47509, EMD-47510, EMD-47511, EMD-47512, and EMD-47513.
- This study does not employ any original code.
- Any additional information required to reanalyze the data reported in this work is available from the lead contact upon request.

### Experimental model and study participant details

The cells used for generating baculovirus are Sf9 cells (*Spodoptera frugiperda,* Cat# CRL-1711), which were cultured at 27°C in suspension, in Sf-900 III SFM (Thermo Fisher Scientific). HEK293S GnTI^-^ suspension cells used for expression of hASIC1a for structural studies were obtained from the ATCC (Cat# CRL-3022) and cultured at 37°C and 8% CO_2_ in a humidified environment. HEK293S GnTI^-^ cells were cultured in FreeStyle 293 expression medium (Thermo Fisher Scientific) supplemented with 2% fetal bovine syndrome (Thermo Fisher Scientific). HEK293T cells were originally purchased from ATCC, Virginia, United States of America. HEK293T cell line in which endogenous hASIC1a was removed by CRISPR/Cas9 (hereafter HEK293T*) was previously described^30^. HEK293T* cells were grown in monolayer in T75 flasks (Orange Scientific) at 37°C in a humidified 5% CO_2_ atmosphere. The culture medium was Dulbecco’s modified Eagle’s medium (DMEM) (Thermo Fisher Scientific) supplemented with 10% fetal bovine serum (Thermo Fisher Scientific) and 1% penicillin-streptomycin (10,000 U/ml; Thermo Fisher Scientific).

## Method details

### Expression and purification of hASIC1a

HEK293S GnTI^-^ cells were grown in suspension to a density of 3.5×10^6^ cells per mL. Cells were infected with baculovirus encoding a full-length human ASIC1a N-terminally tagged with eight histidines (8xHis) and enhanced green fluorescent protein (eGFP), with a thrombin cleavage site between eGFP and hASIC1a (Figure S7A). Cells were incubated for 8-10 hours at 37°C, after which sodium butyrate was added to 10 mM, and the cells transferred to 29°C. For expression of hASIC1a-T26V, 10 μM amiloride was also added to the culture media at this time. After total incubation time of no more than 48 hours post virus addition, the cells were pelleted and pellets washed with Tris-buffered saline before another round of centrifugation. The resultant pellet was snap-frozen in liquid nitrogen and stored at -80℃. Small aliquots of these cells were reserved for preliminary evaluation of GFP-containing protein using fluorescence-detection size-exclusion chromatography (FSEC) (Figure S7B).

The frozen pellet was resuspended in Buffer A (20 mM HEPES pH 7.5, 150 mM NaCl, 0.01 mg/mL deoxyribonuclease I, protease inhibitor cocktail (Pierce)). Cells were lysed by sonication. The lysed cells were centrifuged (8000 x g, 20 min, 4°C), and the supernatant collected. Membranes were harvested by centrifugation (100,000 x g, 1 h, 4°C). The pelleted membranes were snap frozen in liquid nitrogen and stored at -80°C.

The material was re-suspended in fresh Buffer A, homogenized using a Dounce homogenizer, and 1% (w/v) digitonin added. Membrane proteins were solubilized for 1 h at 4℃, then debris removed by centrifugation at 100,000 x g for 1h at 4°C. Homemade GFP nanobody (GNB) affinity resin (1 mL packed resin per g of membrane) was pre-equilibrated in Buffer B (20 mM HEPES pH 7.5, 150 mM NaCl, 0.1% (w/v) digitonin) and transferred to a glass flex-column. Supernatant was then slowly flowed over the column to allow for protein association. The column was washed with 5 column volumes (cv) Buffer B with 2 mM ATP and 2 mM MgCl_2_, then 5 cv Buffer B with nuclease (Pierce), and finally 5 cv of Buffer B with 5 mM CaCl_2_. The resin was then incubated in Buffer B with 5 mM CaCl_2_ and 30 mg thrombin (Prolytix) per mL of resin at room temperature for 1 hour. Cleaved hASIC1a was eluted with 3 cv of Buffer B and collected in 1 cv fractions. Peak fractions were concentrated using a 50 kDa MWCO centrifugal filter (Millipore), and run on a Superose 6 Increase 10/300 gel filtration column (Cytiva) in gel filtration buffer (50 mM sodium phosphate, 200 mM NaCl, 1 mM TCEP, 0.1% (w/v) digitonin). The pH of the buffer varied between 8.5, 7.5, 6.5, and 5.7 depending on the desired final condition of the protein. Peak fractions were pooled and concentrated to ∼4 mg/mL. For certain experiments, MitTx (Alomone Labs) was added at a 1:1 molar ratio (ASIC monomer to toxin). All samples used for grid preparation had fluorinated octyl maltoside (Anatrace) added to a final concentration of 10 μM.

Quantifoil R 2/1 200 mesh Au grids were glow discharged at 15 mA for 30 s. A Vitrobot Mark IV (Thermo Fisher Scientific) was used for blotting and freezing. In general, a 3 µL drop was applied to the surface, then promptly wicked away using a torn piece of filter paper. A second drop of the same volume was applied prior to blotting and plunge freezing in liquid ethane (blot time 3.5 s, wait time 0 s, drain time 0 s, blot force 1, 100% humidity, 12°C). Grids were clipped under vapor-phase nitrogen cooled by liquid nitrogen, before being stored under liquid nitrogen prior to screening and collection.

Protein quality was assessed using FSEC (Figure S7C) and SDS-PAGE (Figure S7D). The latter technique resolved three distinct bands, one of which was of a size consistent with intact hASIC1a monomer. The other two bands were characterized with N-terminal sequencing, and were revealed to be two segments of a cut hASIC1a monomer, most likely a result of off-target cleavage. FSEC traces and cryo-EM maps suggested this partial cleavage in the primary structure did not impact the tertiary structure of the protein (Figures S7D-S7G).

### Image acquisition and processing

Grids were screened on a 200 kV Glacios TEM (Thermo Fisher Scientific) with a K3 Detector (Gatan), and promising candidates underwent data collection on a 300 kV Titan Krios (Thermo Fisher Scientific) with a K3 Detector at the Pacific Northwest Center for Cryo-EM (PNCC) or the HHMI Janelia Research Campus.

Following collection, map processing took place using cryoSPARC^45^ following a generalized workflow. In brief, this included preprocessing with patch motion correction and patch CTF fitting followed by blob picking, several rounds of 2D classification, and *ab initio* map generation. This *ab initio* map was then used to generate templates for extraction of a new particle stack, which was used to generate a new map. This map subsequently underwent multiple rounds of heterogeneous refinement and 3D classification to remove low quality particles from the stack and to evaluate potential conformational heterogeneity. Successive iterations of local refinement, non-uniform refinement^46^, and global and local CTF refinement resulted in the final map (Figure S7H).

For hASIC1a prepared at pH 5.7, multiple structurally unique populations were identified using symmetry expansion prior to focused 3D classification in the transmembrane and micelle region, followed by removal of duplicate particles from each of the three resultant particle stacks. For hASIC1a prepared at pH 7.5, ECD-focused 3D classification yielded two distinct populations, one of which was further classified into two structurally unique populations using heterogeneous refinement (Figure S7H). To enhance TMD map quality and supported by apparent lack of symmetry-breaking structural elements, the maps of C_DS-8.5_, C_DS-7.5_, D_DS-5.7_ and D_RDS-5.7_ were processed with C3 symmetry; all others were processed using C1 symmetry.

### Model building and refinement

Models were built using a combination of Coot^47^ and the Isolde plugin for ChimeraX^48,49^, using a cASIC1 model (PDB ID: 6VTL)^12^ as the starting model. The modeling process consisted of iterative rounds of model building and real-space refinement in Phenix^50,51^. Validation of final models was done using MolProbity^52^. Pore profiles were characterized using HOLE^25^. Model-model positional RMSDs were calculated in PyMol^53^; relative rotational displacements between models were measured using the AngleBetweenHelices plugin for PyMol. Model and map figures were prepared using PyMol^53^ and ChimeraX^48^.

### Whole-cell patch-clamp electrophysiology

For patch-clamp experiments HEK293T* cells were seeded in a T25 Flask (Orange Scientific) and incubated for 24 hours at 37°C before the cells were transfected with 2 µg cDNA coding for hASIC1a (inserted into pcDNA3.1+ vector upstream IRES GFP) using Polyethylenimine (PEI) 25K (Polysciences) in a 1:3 ratio. Cells were used for patch clamp experiments 24 to 36 hours after transfection. On the day of patch clamp recordings, the HEK293T* cells expressing hASIC1a were reseeded on glass coverslips to enable lifting of the cell from the coverslip and positioning in front of a perfusion tool to facilitate rapid solution/pH exchange. The custom-built glass perfusion tool equipped with four adjacent barrels was mounted on a piezo head (PZT head, Siskiyou) and controlled by the MXPZT-300 solution exchanger (Siskiyou). The cells were voltage-clamped at -40 mV at room temperature in the whole-cell configuration using an Axopatch 200B amplifier (Molecular Devices). All recordings were performed using the pCLAMP 10 software (Molecular Devices) and an Axon Digidata 1550A digitizer (Molecular Devices) at 10 kHz. Patch pipettes with a resistance between 3 and 5 MΩ were pulled on a pipette-puller (Model P-1000, Sutter instruments) using borosilicate glass capillaries (World Precision Instruments). The physiological extracellular recording solutions contained 150 mM NaCl, 5 mM KCl, 1 mM MgCl_2_, 2 mM CaCl_2_, 10 mM HEPES, 10 mM D-glucose, and were adjusted to pH 5.6, 6.9 and 8.0 with NaOH and HCl. The phosphate recording solution contained 90 mM NaCl, and 5 mM sodium phosphate. Phosphate buffers of varying pH were created using different ratios of 0.1 M sodium phosphate monobasic and sodium phosphate dibasic solutions, to generate a sodium phosphate stock solution at the desired pH. The intracellular solution consisted of 30 mM NaCl, 120 mM KCl, 1 mM MgCl_2_, 0.5 mM CaCl_2_, 5 mM EGTA, 2 mM Na_2_ATP and 10 mM HEPES adjusted to pH 7.4. pCLAMP10 (Molecular Devices) was used for analyzing current amplitudes and rate constants were fit with single exponential functions to the decay of the current. Graphs were generated using GraphPad Prism 10.

### Two-electrode voltage-clamp electrophysiology and voltage-clamp fluorometry

The human ASIC1a cDNA with a C-terminal 1D4 tag was cloned into a pcDNA3.1+ vector, obtained from TWIST bioscience. Mutations T26V and K105C were introduced using custom-made DNA mutagenesis primers (Eurofins Genomics) and site-directed mutagenesis (PfuUltraII Fusion polymerase, Agilent), followed by sequencing the full coding frame (Eurofins Genomics or Macrogen). Xba1 was used for linearization of cDNAs before transcription to capped cRNA with the Ambion mMESSAGE mMACHINE T7 kit (Thermo Fisher Scientific). *Χenopus* oocyte preparation and the recording setup were the same as previously described^32^. Recording solutions were either Ca^2+^-free ND96 (96 mM NaCl, 2 mM KCl, 1.8 mM BaCl_2_, 1 mM MgCl_2_, 5 mM HEPES) pH adjusted with HCl or NaOH, or Phosphate buffer (90 mM NaCl, 5 mM sodium phosphate, see above). For experiments with MitTx, 0.05% BSA (≥98% essentially fatty acid-free, Sigma-Aldrich) was added to the buffer on the day of use. For SSD experiments, cells were pre- conditioned for 2 min prior to activation with pH 6.0. MitTx was applied for 2 minutes or until a plateau was reached. The reversal potential was established by running a 200 ms voltage-ramp from -60 mV to +60 mV through peak activation at pH 6.5, or during maximal activation by 20 nM MitTx at pH 6.5. For VCF experiments, oocytes expressing hASIC1a K105C were labeled for 30 min in ΟR2 solution (82.5 mM NaCl, 2 mM KCl, 1 mM MgCl_2_, pH 7.4) containing 10 μΜ Alexa Fluor 488 C_5_ maleimide (Thermo Fisher Scientific). Oocytes were then washed twice with OR2 solution and stored in the dark until further use. Recordings and data analysis were done as previously described^32^.

