## Supplementary figures and images for "Conformational plasticity of human acid-sensing ion channel 1a"

### Supplementary figures

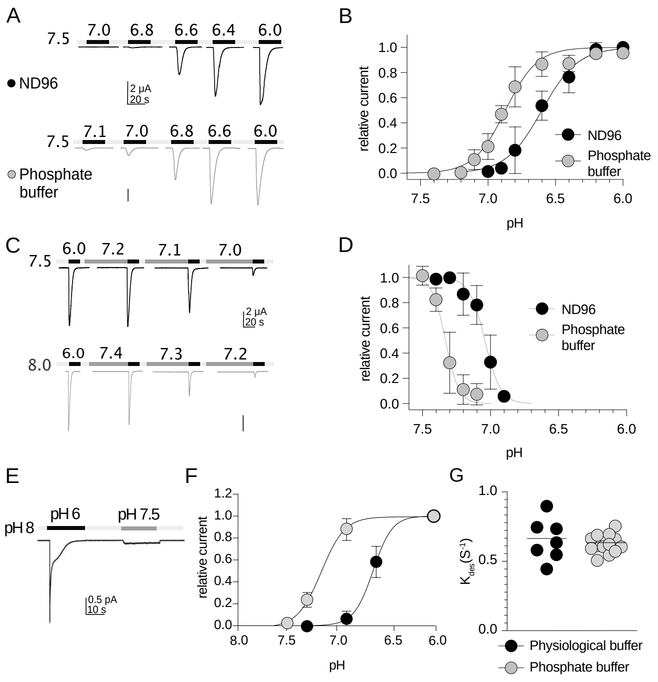

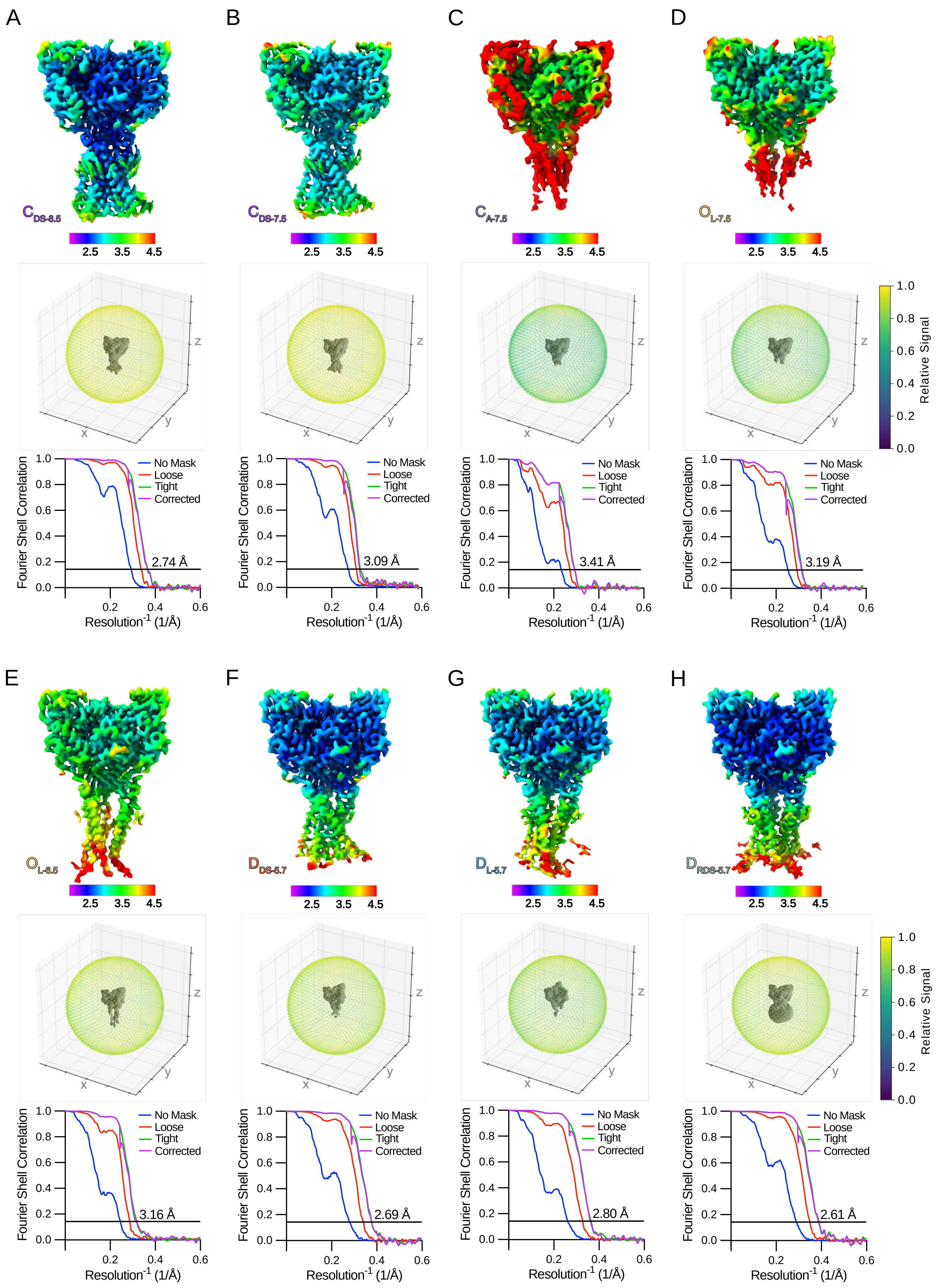

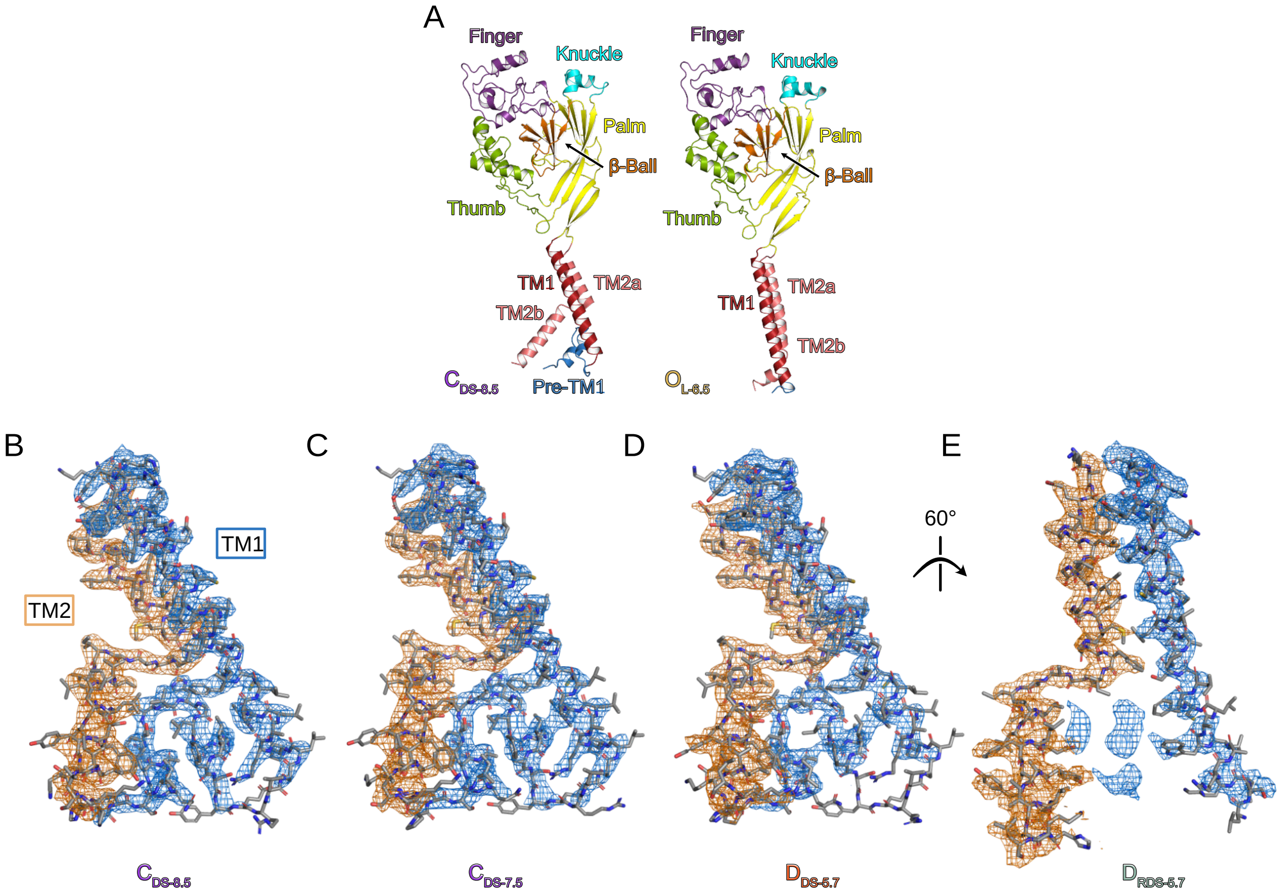

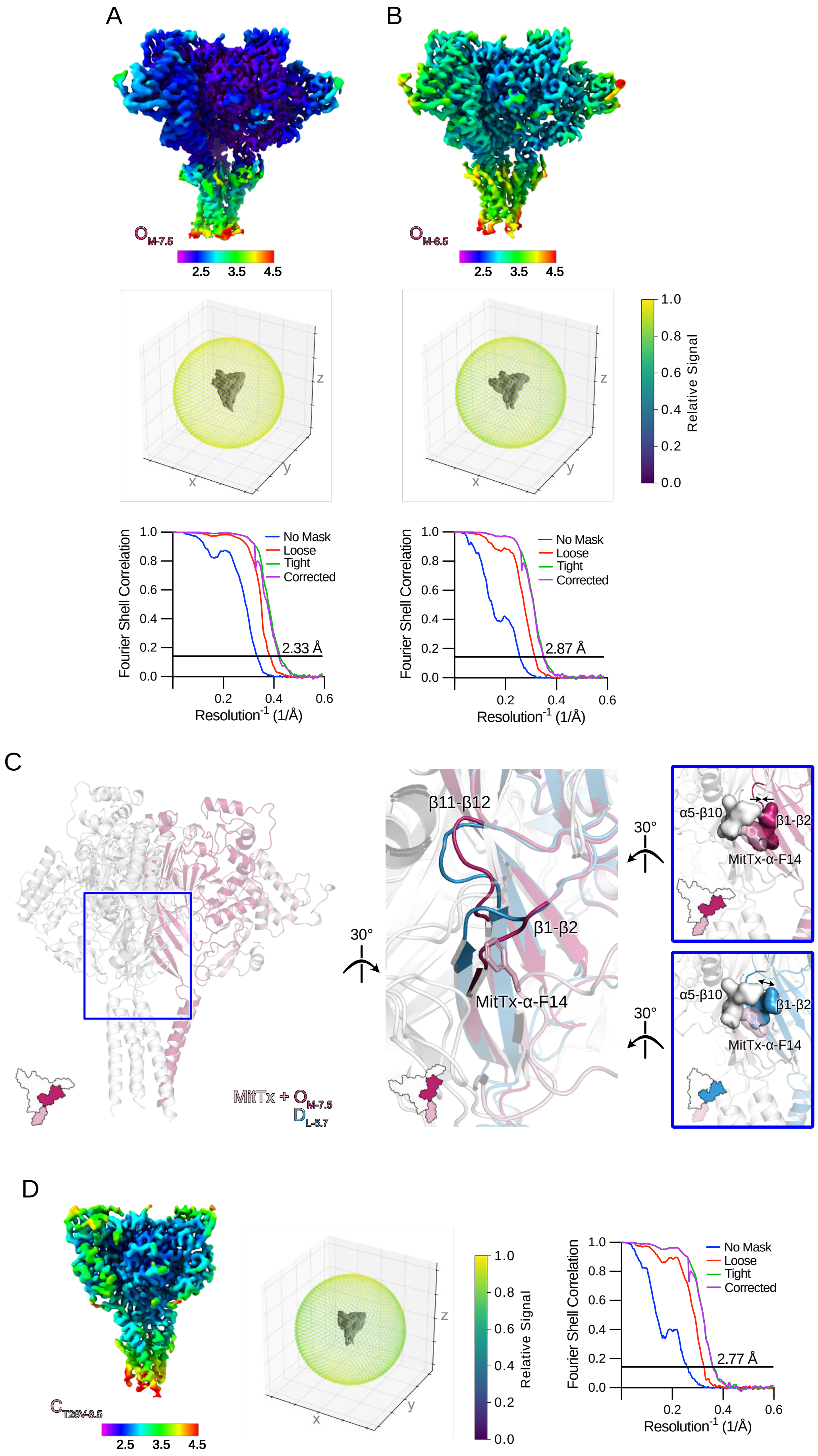

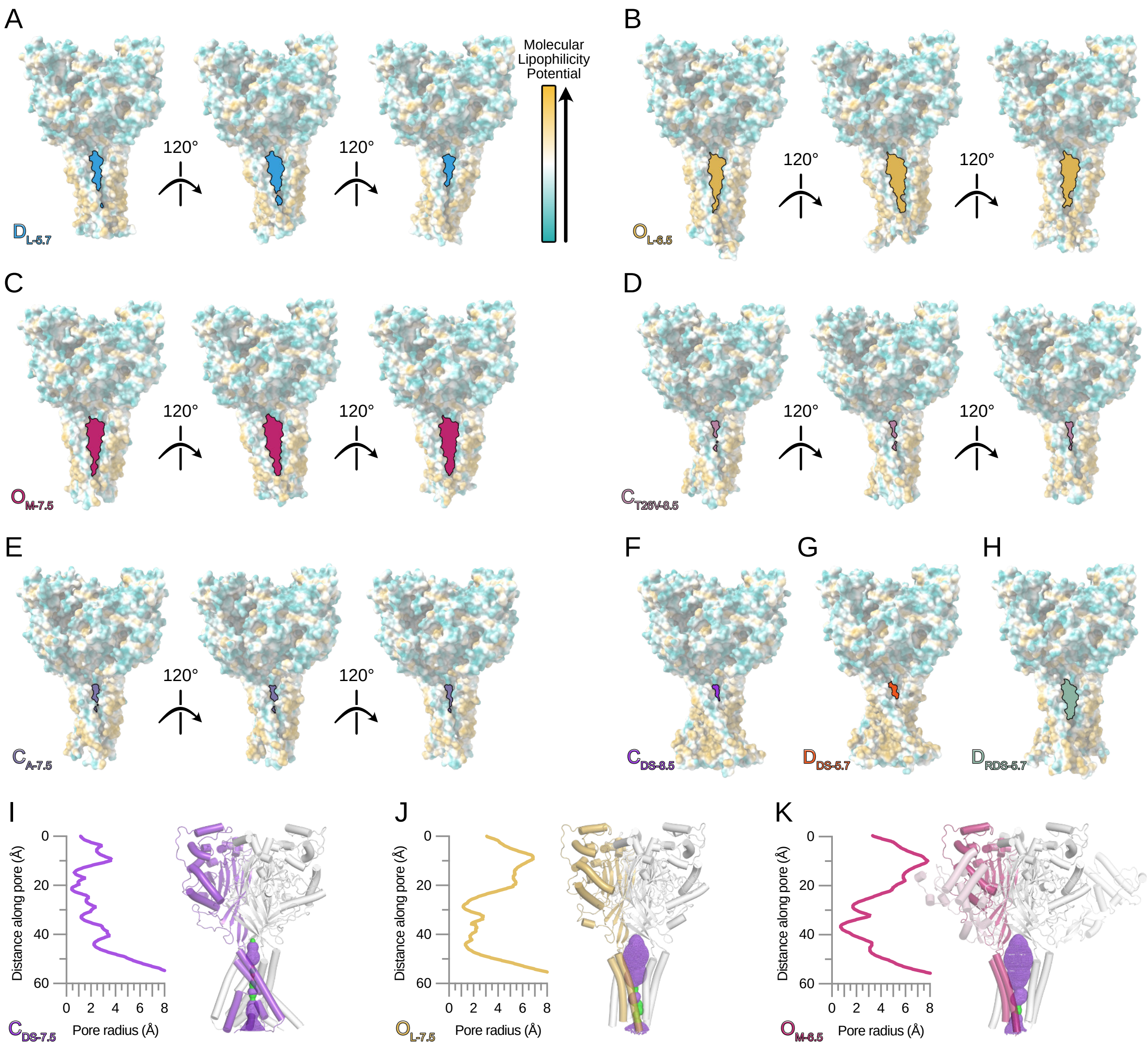

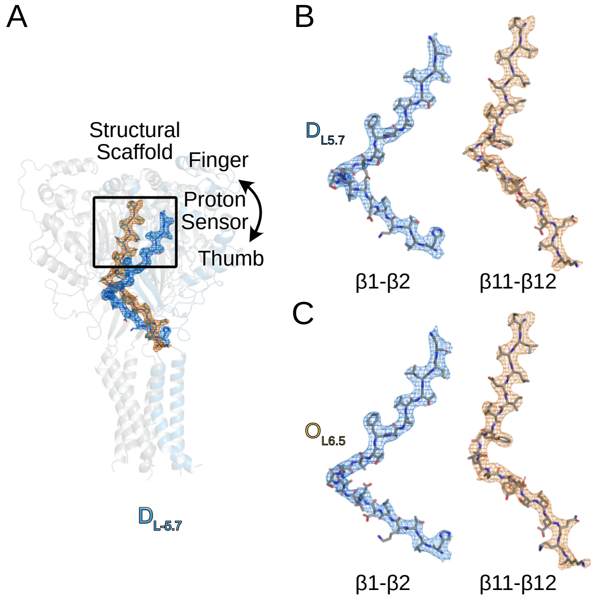

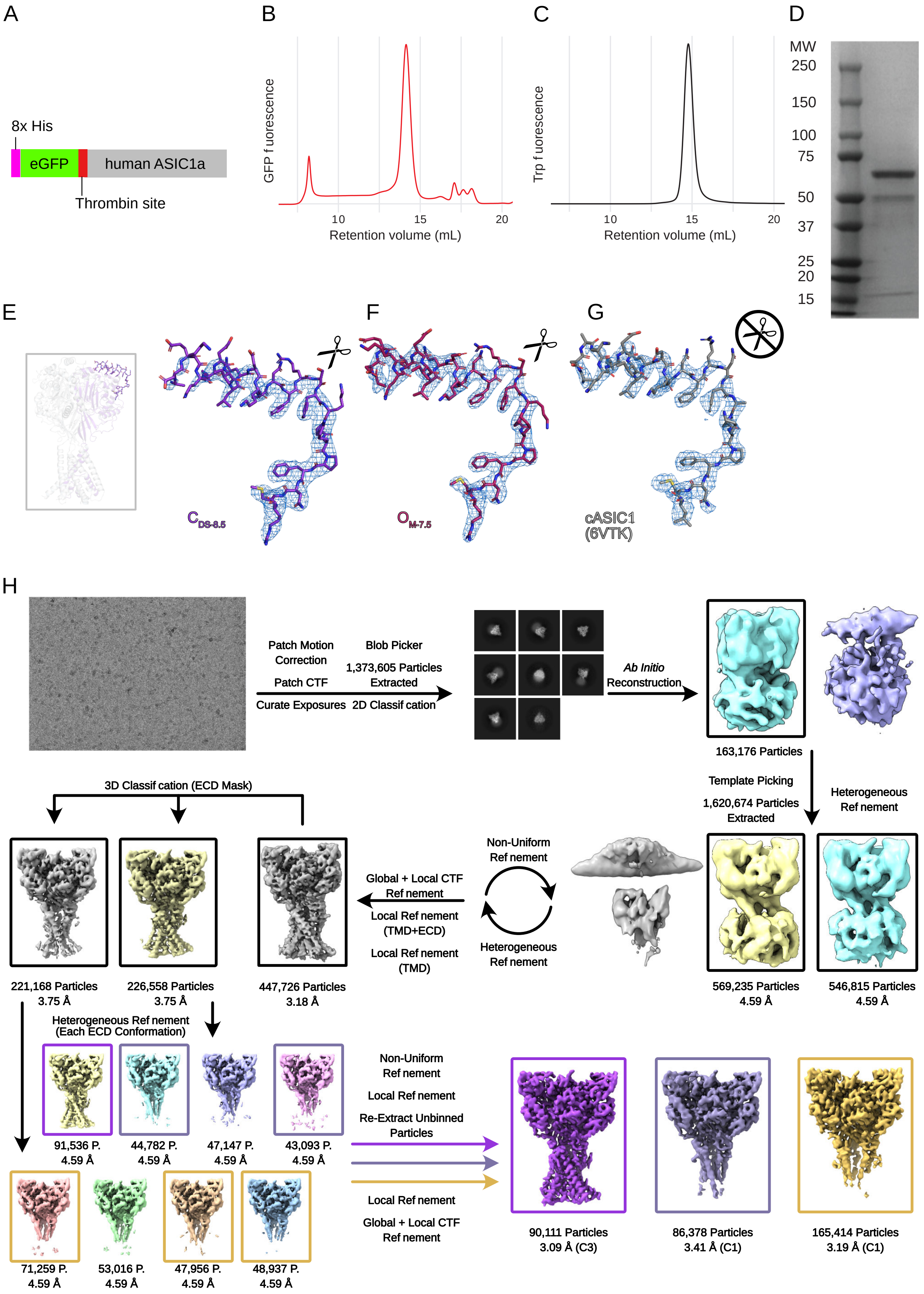
