## Supplementary tables for "Conformational plasticity of human acid-sensing ion channel 1a"

Table S1: Summary of electrophysiology data, related to Figures 2, 4, and S1

| **hASIC1a WT expressed in HEK293T* cells** | **Physiological buffer** | **Phosphate buffer** |
|---|---|---|
| $pH_{50}$ activation | 6.63 ± 0.02 | 7.16 ± 0.03 |
| Desensitization kinetics (rate constants, $S^{-1}$) | 0.66 ± 0.10 (n=7) | 0.63 ± 0.12 (n=13) |
| **hASIC1a WT expressed in *Xenopus* oocytes** | **ND96 (without $Ca^{2+}$)** | **Phosphate buffer** |
| *pH response* | | |
| $pH_{50}$ activation | 6.61 ± 0.05 (n=8) | 6.87 ± 0.05 (n=11) |
| $pH_{50}$ SSD | 7.05 ± 0.07 (n=10) | 7.32 ± 0.04 (n=8) |
| *MitTx response relative to pH 6.0* | | |
| MitTx (20 nM) at pH 8.0 | 0.12 ± 0.10 (n=14) | 0.58 ± 0.10 (n=14) |
| MitTx (20 nM) at pH 6.5 | - | 0.78 ± 0.19 (n=6) |
| MitTx (20 nM) after desensitization at pH 6.5 | - | 0.77 ± 0.12 (n=5) |
| *Reversal potential (mV)* | | |
| Reversal potential at pH 6.5 | 16.92 ± 11.50 (n=11) | 18.00 ± 10.51 (n=13) |
| Reversal potential in MitTx (20nM) at pH 6.5 | -1.17 ± 6.74 (n=9) | -2.80 ± 18.11 (n=9) |
| **hASIC1a K105C expressed in *Xenopus* oocytes** | **Current** | **Fluorescence** |
| *Response relative to pH 6.0* | | |
| pH 7.2 | 0.01 ± 0.02 (n=7) | 0.58 ± 0.16 (n=7) |
| 5 nM MitTx | 0.07 ± 0.02 (n=4) | 0.02 ± 0.02 (n=4) |
| 20 nM MitTx | 0.74 ± 0.17 (n=5) | 0.39 ± 0.09 (n= 5) |
| 50 nM MitTx | 0.77 ± 0.09 (n= 9) | 0.74 ± 0.22 (n= 9) |

Table S2: PDB and EMDB codes of hASIC1a structures, related to Figures 1, 2, and 3

| Name | PDB | EMDB |
|---|---|---|
| $C_{DS-8.5}$ | 9E4A | EMD-47503 |
| $C_{DS-7.5}$ | 9E4B | EMD-47504 |
| $C_{A-7.5}$ | 9E4C | EMD-47505 |
| $O_{L-7.5}$ | 9E4D | EMD-47506 |
| $O_{L-6.5}$ | 9E4E | EMD-47507 |
| $D_{DS-5.7}$ | 9E4F | EMD-47508 |
| $D_{L-5.7}$ | 9E4G | EMD-47509 |
| $D_{RDS-5.7}$ | 9E4H | EMD-47510 |
| $O_{M-7.5}$ | 9E4I | EMD-47511 |
| $O_{M-6.5}$ | 9E4J | EMD-47512 |
| $C_{T26V-8.5}$ | 9E4K | EMD-47513 |

Table S3: Cryo-EM data collection, refinement, and validation statistics, related to Figures 1, 2, and 3

| | $C_{DS-8.5}$ | $C_{DS-7.5}$<br>$C_{A-7.5}$<br>$O_{L-7.5}$ | $O_{L-6.5}$ | $D_{DS-5.7}$<br>$D_{L-5.7}$<br>$D_{RDS-5.7}$ | $O_{M-7.5}$ | $O_{M-6.5}$ | $C_{T26V-8.5}$ |
|---|---|---|---|---|---|---|---|
| Voltage (kV) | 300 | 300 | 300 | 300 | 300 | 300 | 300 |
| Energy filter slit width (eV) | n/a | 20 | n/a | 20 | n/a | 20 | n/a |
| Detector | K3 | K3 | K3 | K3 | K3 | K3 | K3 |
| Operation mode | Non-CDS | CDS | Non-CDS | CDS | Non-CDS | CDS | Non-CDS |
| Total electron exposure on sample ($e^-/Å^2$) | 65 | 65 | 50 | 65 | 50 | 65 | 65 |
| Number of movie frames | 66 | 65 | 45 | 65 | 45 | 65 | 66 |
| Magnification | 105kx | 105kx | 105kx | 105kx | 105kx | 105kx | 105kx |
| Pixel size (Å) | 0.394 | 0.422 | 0.41275 | 0.4195 | 0.41275 | 0.422 | 0.4135 |
| Targeted defocus range (µm) | 0.8-2.2 | 0.8-2.2 | 0.8-2.2 | 0.8-2.2 | 0.8-2.2 | 0.8-2.2 | 0.8-2.2 |
| Number of collected movies | 6540 | 6090 | 7142 | 4600 | 7006 | 4785 | 7621 |
| Symmetry imposed | C3 | C3 ($C_{DS-7.5}$)<br>C1 ($C_{A-7.5}$)<br>C1 ($O_{L-7.5}$) | C1 | C3 ($D_{DS-5.7}$)<br>C1 ($D_{L-5.7}$)<br>C3 ($D_{RDS-5.7}$) | C1 | C1 | C1 |
| Initial particle images (no.) | 855,037 | 1,620,674 | 1,431,341 | 1,610,888 | 1,953,574 | 778,825 | 2,759,695 |
| Final particle images (no.) | 170,179 | 90,111 ($C_{DS-7.5}$)<br>86,378 ($C_{A-7.5}$)<br>165,414 ($O_{L-7.5}$) | 110,918 | 54,398 ($D_{DS-5.7}$)<br>100,628 ($D_{L-5.7}$)<br>84,730 ($D_{RDS-5.7}$) | 541,317 | 100,508 | 102,014 |
| Map resolution (Å) (FSC=0.143) | 2.74 | 3.09 ($C_{DS-7.5}$)<br>3.41 ($C_{A-7.5}$)<br>3.19 ($O_{L-7.5}$) | 3.16 | 2.69 ($D_{DS-5.7}$)<br>2.80 ($D_{L-5.7}$)<br>2.61 ($D_{RDS-5.7}$) | 2.33 | 2.87 | 2.77 |

Table S4: Model refinement and validation statistics, related to Figures 1, 2, and 3

| | $C_{DS-8.5}$ | $C_{DS-7.5}$ | $C_{A-7.5}$ | $O_{L-7.5}$ | $O_{L-6.5}$ | $D_{DS-5.7}$ | $D_{L-5.7}$ | $D_{RDS-5.7}$ | $O_{M-7.5}$ | $O_{M-6.5}$ | $C_{T26V-8.5}$ |
|---|---|---|---|---|---|---|---|---|---|---|---|
| EMDB ID | EMD-47503 | EMD-47504 | EMD-47505 | EMD-47506 | EMD-47507 | EMD-47508 | EMD-47509 | EMD-47510 | EMD-47511 | EMD-47512 | EMD-47513 |
| PDB ID | 9E4A | 9E4B | 9E4C | 9E4D | 9E4E | 9E4F | 9E4G | 9E4H | 9E4I | 9E4J | 9E4K |
| Model resolution (Å) (FSC=0.143) | 2.7 | 3.1 | 3.4 | 3.2 | 3.1 | 2.7 | 2.8 | 2.6 | 2.3 | 2.8 | 2.8 |
| **Model Composition** | | | | | | | | | | | |
| Non-hydrogen atoms | 10842 | 10842 | 9729 | 9915 | 10158 | 10842 | 10024 | 10287 | 14241 | 14285 | 9915 |
| Protein residues | 1353 | 1353 | 1212 | 1239 | 1266 | 1353 | 1251 | 1281 | 1773 | 1778 | 1239 |
| *B* factor ($Å^2$) | 122.72 | 115.31 | 121.13 | 109.71 | 110.83 | 117.56 | 107.11 | 107.97 | 111.44 | 136.75 | 110.98 |
| **R.m.s deviations** | | | | | | | | | | | |
| Bond lengths (Å) | 0.002 | 0.001 | 0.002 | 0.003 | 0.002 | 0.002 | 0.002 | 0.002 | 0.003 | 0.002 | 0.002 |
| Bond angles (°) | 0.460 | 0.394 | 0.542 | 0.442 | 0.414 | 0.451 | 0.501 | 0.478 | 0.451 | 0.428 | 0.420 |
| **Validation** | | | | | | | | | | | |
| MolProbity score | 1.20 | 1.20 | 1.30 | 1.38 | 1.09 | 1.24 | 1.29 | 1.24 | 1.15 | 1.19 | 1.17 |
| Clashscore | 4.16 | 4.16 | 5.54 | 6.96 | 3.00 | 4.72 | 5.37 | 4.68 | 3.59 | 4.05 | 3.84 |
| Poor rotamers (%) | 0.00 | 0.00 | 0.00 | 0.00 | 0.00 | 0.00 | 0.00 | 0.00 | 0.00 | 0.00 | 0.00 |
| **Ramachandran plot** | | | | | | | | | | | |
| Favored (%) | 99.33 | 99.26 | 98.18 | 98.54 | 98.97 | 98.44 | 98.15 | 99.06 | 99.09 | 98.80 | 99.03 |
| Allowed (%) | 0.67 | 0.74 | 1.82 | 1.46 | 1.03 | 1.56 | 1.85 | 0.94 | 0.91 | 1.20 | 0.97 |
| Disallowed (%) | 0.00 | 0.00 | 0.00 | 0.00 | 0.00 | 0.00 | 0.00 | 0.00 | 0.00 | 0.00 | 0.00 |
