## Supplementary material for "Conformational plasticity of human acid-sensing ion channel 1a": Key resources table

### STAR Methods

#### Key resources table

| REAGENT or RESOURCE | SOURCE | IDENTIFIER |
|---|---|---|
| Bacterial and virus strains | | |
| DH10Bac Competent Cells | Invitrogen | Cat# 10361012 |
| XL1-Blue competent cells | Agilent | Cat# 200249 |
| Chemicals, peptides, and recombinant proteins | | |
| Amiloride hydrochloride hydrate | Sigma | Cat# A7410 |
| Human alpha-Thrombin | Prolytix | Cat# HCT-0020 |
| MitTx | Alomone Labs | Cat# M-100 |
| Digitonin | Millipore | Cat# 300410 |
| Octyl Maltoside, Fluorinated, Anagrade | Anatrace | O310F |
| Pierce™ Protease Inhibitor Tablets, EDTA-free | Thermo Fisher Scientific | Cat# A32965 |
| Pierce™ Universal Nuclease for Cell Lysis | Thermo Fisher Scientific | Cat# 88702 |
| Fetal Bovine Serum | Thermo Fisher Scientific | Cat# 10082147 |
| Sf-900™ III SFM | Thermo Fisher Scientific | Cat# 12658027 |
| FreeStyle™ 293 Expression Medium | Thermo Fisher Scientific | Cat# 12338018 |
| Dulbecco's Modified Eagle Medium (DMEM) | Thermo Fisher Scientific | Cat# 41966052 |
| Penicillin-Streptomycin | Thermo Fisher Scientific | Cat# 15140-122 |
| PfuUltraII Fusion polymerase | Agilent Technologies | Cat # 600385 |
| Custom-made DNA mutagenesis primers | Eurofins Genomics | N/A |
| Bovine serum albumin ≥98% essentially fatty acid-free | Sigma-Aldrich | Cat # A9418 |
| Critical commercial assays | | |

| | | |
|---|---|---|
| PureLink™ HiPure Plasmid Miniprep Kit | Invitrogen | Cat# K210002 |
| Superose 6 Increase 10/300 GL | Cytiva | Cat# 29091596 |
| Ambion mMESSAGE mMACHINE T7 kit | Thermo Fisher Scientific | Cat# AM1344 |
| Plasmid DNA purification kit | Geneaid | Cat # PD300 |
| Deposited data | | |
| Coordinates of hASIC1a at pH 8.5 ($C_{DS}$) | This paper | PDB: 9E4A |
| Cryo-EM map of hASIC1a at pH 8.5 ($C_{DS}$) | This paper | EMDB: EMD-47503 |
| Coordinates of hASIC1a at pH 7.5 ($C_{DS}$) | This paper | PDB: 9E4B |
| Cryo-EM map of hASIC1a at pH 7.5 ($C_{DS}$) | This paper | EMDB: EMD-47504 |
| Coordinates of hASIC1a at pH 7.5 ($C_A$) | This paper | PDB: 9E4C |
| Cryo-EM map of hASIC1a at pH 7.5 ($C_A$) | This paper | EMDB: EMD-47505 |
| Coordinates of hASIC1a at pH 7.5 ($O_L$) | This paper | PDB: 9E4D |
| Cryo-EM map of hASIC1a at pH 7.5 ($O_L$) | This paper | EMDB: EMD-47506 |
| Coordinates of hASIC1a at pH 6.5 ($O_L$) | This paper | PDB: 9E4E |
| Cryo-EM map of hASIC1a at pH 6.5 ($O_L$) | This paper | EMDB: EMD-47507 |
| Coordinates of hASIC1a at pH 5.7 ($D_{DS}$) | This paper | PDB: 9E4F |
| Cryo-EM map of hASIC1a at pH 5.7 ($D_{DS}$) | This paper | EMDB: EMD-47508 |
| Coordinates of hASIC1a at pH 5.7 ($D_L$) | This paper | PDB: 9E4G |

| | | |
|---|---|---|
| Cryo-EM map of hASIC1a at pH 5.7 ($D_L$) | This paper | EMDB: EMD-47509 |
| Coordinates of hASIC1a at pH 5.7 ($D_{RDS}$) | This paper | PDB: 9E4H |
| Cryo-EM map of hASIC1a at pH 5.7 ($D_{RDS}$) | This paper | EMDB: EMD-47510 |
| Coordinates of hASIC1a and MitTx at pH 7.5 ($O_M$) | This paper | PDB: 9E4I |
| Cryo-EM map of hASIC1a and MitTx at pH 7.5 ($O_M$) | This paper | EMDB: EMD-47511 |
| Coordinates of hASIC1a and MitTx at pH 6.5 ($O_M$) | This paper | PDB: 9E4J |
| Cryo-EM map of hASIC1a and MitTx at pH 6.5 ($O_M$) | This paper | EMDB: EMD-47512 |
| Coordinates of hASIC1a-T26V at pH 8.5 ($C_{T26V}$) | This paper | PDB: 9E4K |
| Cryo-EM map of hASIC1a-T26V at pH 8.5 ($C_{T26V}$) | This paper | EMDB: EMD-47513 |
| Experimental models: Cell lines | | |
| HEK293S GnTI- | ATCC | Cat # CRL-3022 |
| Sf9 (*Spodoptera frugiperda*) | ATCC | Cat# CRL-1711 |
| HEK293T ASIC1a knockout | ATCC | Originally Cat# CRL-3216 |
| *Xenopus Laevis* oocytes | Xenopus1, US | N/A |
| Recombinant DNA | | |
| Plasmid: pEG BacMam | Gift from Eric Gouaux | doi: 10.1038/nprot.2014.173 |
| pEG-eGFP-hASIC1a | This paper | N/A |
| pEG-eGFP-hASIC1a-T26V | This paper | N/A |
| hASIC1a-1D4 in pcDNA3.1+ vector | Twist Bioscience | N/A |
| hASIC1a-1D4-IRES-eGFP in pcDNA3.1+ | GeneArt, Thermo Fisher Scientific | N/A |
| Software and algorithms | | |

| | | |
|---|---|---|
| CryoSPARC | Doi:10.1038/nmeth.4169 | RRID:SCR_016501 |
| Pymol | Pymol Molecular Graphics System, Schrodinger, LLC | RRID:SCR_000305 |
| Coot | https://doi.org/10.1107/S0907444904019158 | RRID:SCR_014222 |
| UCSF ChimeraX | Doi:10.1002/pro.3235 | RRID:SCR_015872 |
| MolProbity | Doi:10.1107/S0907444909042073 | RRID:SCR_014226 |
| Serial EM | Doi:10.1016/j.jsb.2005.07.007 | http://bio3d.colorado.edu/SerialEM |
| Phenix | Doi:10.1107/S2059798318006551 | RRID:SCR_014224 |
| ISOLDE | Doi:10.1107/S2059798318002425 | https://isolde.cimr.cam.ac.uk/ |
| pClamp 10.5.1.0 | Molecular Devices | RRID:SCR_011323 |
| GraphPad Prism 10.3.1. | GraphPad | RRID:SCR_002798 |
| Other | | |
| Quantifoil Holey Carbon Grids, 2/1 Au 200 mesh grids | SPI Supplies | Cat# 4320G |
